# SUMO interacting motif (SIM) of S100A1 is critical for S100A1 post-translational protein stability

**DOI:** 10.1101/2023.01.18.524665

**Authors:** Zegeye H. Jebessa, Manuel Glaser, Jemmy Zhao, Andrea Schneider, Ramkumar Seenivasan, Martin Busch, Julia Ritterhoff, Rebecca C. Wade, Patrick Most

## Abstract

S100A1 is a small EF-type Ca^2+^ sensor protein that belongs to the multigenic S100 protein family. It is abundantly expressed in cardiomyocytes (CMs) and has been described as a key regulator of CM performance due to its unique ability to interact with structural contractile proteins, regulators of cardiac Ca^2+^ cycling, and mitochondrial proteins. However, our understanding of the molecular mechanisms regulating S100A1 protein levels is limited. We used the bioinformatics tool GPS-SUMO2.0 to identify a putative SUMO interacting motif (SIM) on S100A1. Consistently, a S100A1:SUMO interaction assay showed a Ca^2+^-dependent interaction of S100A1 with SUMO proteins. In neonatal rat ventricular myocytes (NRVM) and COS1 cells, S100A1 protein abundance increased in the presence of overexpressed SUMO1 without affecting the S100A1 mRNA transcript. We then generated S100A1 truncation mutants, where the SIM motif was removed by truncation or in which the core residues of the SIM motif (residues 77-79) were deleted or replaced by alanine. In COS1 cells and NRVM, overexpression of these S100A1 mutants led to elevated S100A1 mutant mRNA levels but failed to produce respective protein levels. Protein expression of these mutants could be rescued from degradation by addition of the proteasome inhibitor MG-132. By using an information-driven approach to dock the three-dimensional structures of S100A1 and SUMO, we predict a novel interaction mode between the SIM in S100A1 and SUMO. This study shows an important role of SUMO:SIM-mediated protein:protein interaction in the regulation of post-translational protein stability, and provides mechanistic insights into the indispensability of the core SIM for S100A1 post-translational stability.

## Introduction

S100 proteins belong to a multigenic protein family. They are characterized by two calcium (Ca^2+^) binding EF-hand domains and are exclusively expressed in vertebrates in a cell-type specific manner. S100 proteins integrate Ca^2+^ signals to regulate pleiotropic, yet specialized cellular functions, such as cell proliferation, differentiation, motility and survival [1-4].

S100A1, a prominent member of the S100 protein family, is predominantly expressed in the myocardium and has emerged as a central regulator of cardiomyocyte (CM) Ca^2+^ homeostasis and cardiac contraction [5-7]. S100A1 expression is downregulated in left-heart failure with reduced ejection fraction (HFrEF) and right-sided (cor pulmonale) heart failure, which contributes to progression and mortality in both diseases [8-11]. Gene regulation analysis of S100A1 suggested a role for transcriptional mechanisms that regulate S100A1 expression [8, 12, 13], but post-translational mechanisms that could regulate protein stability have not been examined yet.

The 3-dimensional structure of S100A1 has been extensively studied by X-ray crystallography and NMR spectroscopy, that allows to comprehend its structure-function relationship [14-20]: S100A1 forms an anti-parallel, symmetric homodimer independently of its Ca^2+^ binding capabilities. The secondary structure of S100A1 consists of four α-helices (H-I, H-II, H-III and H-IV), two loops (L-1 and L-2) and a hinge region (flanked by H-II and H-III). Its two EF-hands are characterized by a 14-residue-long N-terminal pseudo-Ca^2+^-binding domain (located at L-1) and a 12-residue-long C-terminal canonical Ca^2+^-binding domain (located at L-2). The hinge region and the C-terminal extension (downstream of the C-terminal Ca^2+^-binding domain) display the most variation between individual S100 proteins, which has been linked to their specific biological activities. Upon Ca^2+^-binding, the C-terminal extension displays a conformational change and thereby exposes hydrophobic residues, which have been implicated in target protein interaction [1, 21-26]. S100A1 can be modified by several post-translational modifications (PTM) [16, 19, 27] that may not only regulate its activity but also protein stability.

PTM by Small Ubiquitin-like Modifier (SUMO) controls protein function by regulating protein-protein interactions, subcellular localization, or opposing degradation by ubiquitination [28, 29]. For this, SUMO proteins can undergo multivalent interactions: It can be attached to target proteins at exposed lysine residues through SUMOylation, or undergo non-covalent interactions by binding to the SUMO interaction motif (SIM) [29-33]. S100A4, a member of the S100 protein family, can be SUMOylated [34] and several other S100 protein family members were identified in a SUMO:SIM interaction screen to interact with either SUMO1 or SUMO2 [35]. However, whether S100A1 is modified by SUMO proteins has so far not been explored.

We therefore hypothesized that post-translational SUMO-dependent signaling may impact S100A1 protein stability. We used a web-based prediction algorithm and identified a SIM in the S100A1 C-terminal extension on H-IV. Experimentally, S100A1 interacted in a Ca^2+^ dependent manner with SUMO 1 and 2. We identified the residues -_77_VLV_79_-, which constitute the core SIM motif on S100A1, as critical for S100A1 post-translational stability. Finally, computational molecular docking identified a novel S100A1:SUMO interaction mode. Together, these data reveal a yet unrecognized post-translational molecular checkpoint for S100A1’s protein stability involving a novel SIM-mediated interaction between the S100A1 homodimer and SUMO1.

## Experimental procedures

### Experimental Animals

All experimental procedures were in line with the principles of the Declaration of Helsinki. Approval was granted by the Institutional Animal Care and Use Committee at the Regierungspräsidium Karlsruhe, Germany and were performed in accordance with institutional guidelines of the University of Heidelberg.

### GPS-SUMO *In silico* prediction of SUMOylation sites and SUMO binding Motifs (SIM) on S100A1

The human S100A1 sequence in FASTA format (Uniprot ID= P23297) was submitted to the GPS-SUMO 2.0 Online service webserver (http://sumosp.biocuckoo.org/online.php) [36]. The default setting of the webserver was modified and the threshold set to “high” to predict SUMOylation and SUMO interaction. GPS = generation group-based prediction system.

### *In vitro* SUMO Interaction Assay (SIM binding assay)

Bead preparation and *in vitro* binding assays were performed as described previously [37]. In summary, SUMO1 and 2, ubiquitin and ovalbumin proteins were coupled with pre-treated cyanogen bromide-activated sepharose beads (1 g/3.5 ml final bed volume) in NaHCO_3_ buffer pH 8.9 for 3 hours at room temperature (RT). Beads were then washed with NaHCO_3_ buffer and blocked with NaHCO^3+^ 100 mM Tris pH 8.8. Subsequently, the beads were washed with SAB- (20 mM HEPES pH 7.3, 110 mM K-acetate, 2 mM Mg-acetate, 1 mM EGTA, 0.05% Tween20, protease inhibitors (Aprotinin/Leupeptin/Pepstatin 1ug/ml), 1 mM DTT) buffer and blocked with SAB-buffer with 0.2mg/ml ovalbumin. To examine whether purified recombinant S100A1 [38] interacts with SUMO proteins, the SUMO-1, SUMO-2 and ubiquitin carrying sepharose beads as well as the ovalbumin blocked sepharose beads were pretreated with TBC-buffer (20 mM HEPES pH 7.3, 110 mM K-acetate, 2 mM Mg-acetate, 1 mM CaCl2, 0.05% Tween20, protease inhibitors (Aprotinin/Leupeptin/Pepstatin 1ug/ml), 1 mM DTT) and 10 µg of protein of interest (S100A1) is diluted in TBC+ (0.2 mg/ml Ovalbumin) buffer and with the prepared beads. The beads were washed 3X times with TBC-buffer and 20 µl of sample buffer (loading dye) is added to the beads and incubated for 10 min at 30 °C. The supernatant is eluted and boiled at 96 °C for 5 mins and loaded into PAGE gel analysis and subsequently for imunoblotting.

### *In vitro* SUMOylation Assay

To test whether purified recombinant S100A1 is SUMOylated, an *in vitro* SUMOylation assay was performed according to the manufacturer’s instructions for the SUMOylation assay kit (#ab139470). We modified the assay adding as indicated 2 mM CaCl_2_ or 2 mM EGTA to the SUMOylation reaction buffer.

### Expression plasmids and adenoviruses

WT S100A1 and its variants were synthesized (Eurofins MWG) and cloned into the pAdTrack-CMV plasmid (#16405, Addgene) (Table S1). pAdTrack-CMV plasmid contains two independent CMV promoters, enforcing the expression of the gene of interest and the reporter EGFP (enhanced green fluorescence protein) (Fig S3A). The pAdTrack-CMV were used as an expression vector and as a shuttle vector for the generation of adenoviral vectors.

The N-terminal FLAG-SUMO1 plasmid was received from late Jeffrey Robbins as described previously [39]. For Adenovirus production of Ad.S100A1, Ad.MYC-S100A1, Ad.S100A1-1-74 and Ad.MYC-S100A1-1-74; we employed the pADEasy system of adenovirus production [40]. Briefly, to produce adenovirus recombinants, pAdEasy-1 carrying BJ5183-AD-1 electroporation competent cells (#200157, Agilent technologies) were transformed with *PmeI* linearized pAdTrack-CMV with the gene of interest. The recombinant transformants were selected on kanamycin plates. Plasmid DNAs of positive clones were propagated and purified using the standard Qiagen maxiprep plasmid purification kit (#12163, Qiagen). HEK-293 cells (ATCC type) were transfected with *PacI* linearized recombinant pAdEasy-1 using Lipofectamine 2000 (#11668027, Thermofischer) and subsequently kept in culture for up to 20 days before harvesting and CsCl gradient adenovirus purification [40].

### Cell culture

COS1 cells and HEK293 cells were grown in Dulbecco’s Modified Eagle’s Medium (DMEM) supplemented with fetal calf serum (FCS [10%]), l-glutamine (1%) and penicillin/streptomycin (1%) (complete DMEM).

HEK293 cells were used for generation, production and propagation of adenoviruses used in the experiment. COS1 cells were transfected with pAdTrack-CMV expression plasmids (Table S1) using the GeneJammer transfection reagent (#204130, Agilent Technologies) according to the manufacturer’s protocol. Wherever indicated, 38 hours post transfection, COS1 cells were treated with 10 µM of MG-132 (#M7449-200UL, Sigma Aldrich) for 10 hours before harvest.

### NRVM isolation and culture

NRVMs were isolated from 1- to 3-day-old Wistar-rats (Janvier) following standard enzymatic digestion and percoll gradient purification procedures [41]. Briefly, hearts from neonatal rats were rapidly excised and washed in sterile PBS to remove blood and debris. Next, the ventricles were minced and dissociated into single cells by repeated sequential pancreatin (Sigma, [#8049-47-6, lot#SLBN2032V]) and collagenase II (Worthington, [#LS004177, lot#S5B15572]) digestion (37°C) with gentle stirring. The enzymatic digestion was inactivated by adding FCS and removed from dissociated cells by centrifugation (1000 rpm, RT) and the eventual pellet was resuspended in FCS. Before NRVM purification in ADS-percoll gradient, the sequentially digested NRVMs were pooled, centrifuged and the pellets were resuspended in ADS buffer (116.4 mM NaCl, 19.7 mM HEPES, 9.42 mM NaH_2_PO_4_ x H_2_O, 5.55 mM Anhydrous-Glucose, 5.36 KCl, 0.83 Anhydrous-MgSO_4_, pH7.4). To purify NRVMs; the percoll (#17-0891-01, GE healthcare) gradient was prepared using ADS buffer. The bottom and top layers of ADS-percoll were made and NRVMs in ADS were added, and subsequently centrifuged at 2400 rpm, 4°C for 30 minutes. The purified NRVM layer was then plated in complete DMEM in a cell culture dish. The next day, NRVMs were transduced using adenoviruses (10 M.O.I). In experiments involving proteasomal inhibition, NRVMS were treated with proteasomal inhibitor MG-132 (#M7449-200UL, Sigma Aldrich) for 10 hours prior to harvest.

### ARVM isolation and culture

Adult rat ventricular myocytes (ARVM) were isolated following standard enzymatic digestion and as described previously [41, 42]. ARVMs were then maintained in M199 supplemented with taurine (5 mM), carnitine (5 mM), creatine (4.4 mM), penicillin/streptomycin (1%), Insulin/Insuman® (Sanofi, 25 I.U), Mercaptopropionylglycine (4.9 mM). 2 hours after plating, ARVMs were then transduced with adenovirus at 10 MOI 24 hrs. For the immunofluorescence assay, ARVMs were treated with 10µM of MG-132 for 10 hours prior to fixation in 4% paraformaldehyde (PFA).

### Immunoblotting/Western blotting

COS1 cells and NRVM were harvested and lysed in cytolysis buffer with protease and phosphatase inhibitors (10 mM HEPES pH 7.9, 1.5 mM MgCl_2_, 10 mM KCl, 0.5 mM DTT, 0.05% NP40, protease inhibitor cocktail tablets (Roche) and cocktail-2 and cocktail-3 phosphatase inhibitors (Sigma Aldrich)). All samples were then analyzed by SDS-PAGE (self-made 16.5% SDS-PAGE) and transferred onto a PVDF membrane before probing with the indicated antibodies. The following antibodies were used: anti-S100A1 (#ab11428, abcam), anti-S100A1 (#SP5355P, acris), anti-GFP (#2956, cell signaling), anti-FLAG (#F1804, sigma) and anti-MYC (#2276, cell signaling) and were used at 1:1000 dilution; anti-GAPDH was obtained from EMD Millipore (#MAB374) and was used at 1:10,000 dilution, anti-SUMO1 and anti-SUMO2/3 were obtained from abcam together with the SUMOylation assay kit (#ab139470) and were used at 1:1000 dilution. Goat anti-mouse (IRDye 680RD, #925-68070 or IRDye 800CW, #926-32210) and goat anti-rabbit (IRDye 680, #926-68071 or IRDye 800CW, #926-32211) secondary antibodies were used at 1:5000 and obtained from Li-cor. Signal intensity quantification analysis was conducted using Image studio light version 5.2 (Li-cor).

### Immunostaining (Indirect immunofluorescence)

ARVM were plated at 50,000 cells/well on laminin coated (#L2020, Sigma-Aldrich) 4 well chamber slides (#94.6150.401, Sarstedt) in 1 ml M199 complete medium and transduced, as indicated, with adenovirus. ARVM were then fixed with 4% paraformaldehyde (PFA) in PBS before being permeabilized with Triton X-100 (0.1%), blocked with 1 % BSA in PBS and stained with antibody against anti-MYC (#2276, cell signaling) at 1:1000 and Isotype control Mouse IgG2a (#14-4724-82, ThermoFisher scientific) at 1:1000. Alexa Fluor 546 anti-mouse secondary antibody (A10036, ThermoFisher scientific) at 1:1000 was used to detect MYC signal. The ARVM were counter stained with DAPI at 1:3000 in PBS for 10 minutes.

### Gene expression assay

Total RNA was isolated and purified from cultured cells using a standard commercial RNA prep kit (#48500, Norgen) and 1 μg of RNA was reverse transcribed to cDNA using a cDNA synthesis kit (#205314, Qiagen). 10 ng of cDNA was used for quantitative PCR by SYBR Green. End point PCR to detect expression of the transgene was performed using the Phusion high fidelity polymerase (#M0530L, NEB) protocol. The primers used for quantitative RT-PCR and for end point PCR analysis are shown in Table S2.

### Statistics and Reproducibility

All experimental data are expressed as mean ± SEM (Standard Error of Mean). N-numbers are given in the figure legends. Statistical analysis was performed using GraphPad prism 9 (GraphPad Software, San Diego, California, USA). An unpaired two-tailed t-test was used to compare two groups. An ordinary one-way ANOVA with Bonferroni’s post-hoc multiple comparison test was used to compare three and more groups. For all comparison, a p < 0.05 was considered significant and exact p values are given in the figures.

### Modeling and analysis of SUMO1-S100A1 complexes

Docking of the proteins was performed using the HADDOCK 2.4 webserver [43, 44], with default settings except that the option for final refinement by short molecular dynamics simulations in explicit solvent was switched on to optimize the docking poses. The structure of SUMO1 used for docking was extracted from the crystal structure with PDB id 6V7Q [45] (*Homo sapiens*, C52A mutant; bound to PIAS-derived phosphorylated peptide containing SIM; residues 20-95, chain A, alternate location A for R54). The NMR structure with PDB id 2LP3 [46] (*Homo sapiens*; residues 2-94, chain A/B) of the Ca^2+^-bound wild-type S100A1 homodimeric complex was chosen as the input structure for holo-docking, while the NMR structure with PDB id 2L0P [47] (*Homo sapiens*; residues 2-94, chain A/B) of the Ca^2+^-unbound wild-type S100A1 homodimeric complex was selected as the input structure for the apo-dockings. The UniProt-numbering is used throughout the text. Clustering of the 20 2LP3 or 2L0P NMR structural models was performed with cpptraj [48] from the Amber18 molecular dynamics package [49], applying the average linkage hierarchical agglomerative clustering approach together with the root-mean square deviation between the S100A1 Cα atoms as a distance criterion. The number of target clusters was set to five. Docking was then performed separately to the different cluster representatives. To specify the ambiguous interaction restraints used by HADDOCK for docking, the S100A1 residues corresponding to the interacting residues in the SIM predicted by GPS-SUMO2.0 in the S100A1 H-IV were defined as active residues in the input form (i.e., _76_VVLVA_80_ in each subunit/chain of the S100A1 homodimer). For docking of SUMO1 to the S100A1-AAA mutant, the _76_VAAAA_80_ residues were defined as active residues in the input form. Residue mutations were handled by HADDOCK (after manually deleting side-chain coordinates and renaming residues accordingly). For SUMO1, two sets of active residues were probed: (i) set I corresponds to the SUMO1 residues 30-55 (i.e., the groove formed by the α1-helix and the β2-strand of SUMO1 plus the loop connecting them; see below) and (ii) set II corresponds to the SUMO1 residues 28, 29, 63, 66-70, 75, 81, 83, 86, 89, 91 (relevant residues located on the opposite side of the groove formed by the α1-helix and the β2-strand; see below).

Binding affinity was estimated by using two conceptually different approaches: (i) the ‘AnalyseComplex’-functionality of the FoldX [50] 5.0 Suite command-line tool, which estimates affinities by computing the difference between bound and unbound states using the FoldX force field, including terms that are relevant for protein stability, and (ii) Prodigy [51], which is a linear model that predicts protein-protein binding affinities from intermolecular contacts in the complexes (but does not consider the Ca^2+^ ions). As recommended, the structures were additionally “repaired” using the ‘RepairPDB’-function of the FoldX 5.0 command-line tool before running FoldX 5.0 binding estimation computations. The S100A1-SUMO1 binding affinities were estimated using the final docked poses, averaging over the four top-scoring poses from the best-scoring clusters, as the four top-scoring poses were also employed for the HADDOCK cluster scoring. Prodigy was also used to obtain intermolecular residue contacts. Polar contact analysis and structural superpositions were performed with PyMOL Version 2.3.0 [52].

Brownian dynamics-based dockings were performed with webSDA 1.0 [53], which runs the SDA (Simulation of Diffusional Association) 7.2.2 software [54-57], using default settings if not specified otherwise. No prior knowledge about potentially interacting residues was used, with the condition to record an encounter complex being solely the closeness of atoms representative of the centers of geometry of the docking partners (the maximum distance between the corresponding atoms to record a complex was the default value of 54.8995 Å). Ca^2+^ ions, assigned a charge of +2e and a van der Waals’ radius of 1.7131 Å, were inserted manually in webSDA-generated PQR files[58] in order to incorporate them as effective charge sites. Electrostatic interaction [59, 60], electrostatic desolvation [61], and non-polar desolvation [62] terms were used to compute intermolecular forces. 5000 SDA trajectories were generated and up to 2000 encounter complexes were recorded. For the clustering of encounter complexes, 10 clusters were requested. Electrostatic potentials were computed and visualized with the APBS electrostatics plugin in PyMOL 2.5.2 [58, 59, 63]. Visualization was performed with PyMOL (2.3.0 and 2.5.2) and VMD 1.9.4a55 [64].

## Results

### S100A1 has a SIM and interacts with SUMO proteins

To address the question whether S100A1 can be modified by SUMO, we used the web server-based tool GPS-SUMO 2.0, that allows the prediction of SUMOylation sites and SUMO Interaction Motifs (SIM) [36, 65]. As S100A1 displays high sequence homology between different species, we used human S100A1 (Uniprot ID: P23297) as input sequence. The algorithm identified a SIM at residues 69-87 of human S100A1 containing the core SIM residues 76-80 (_76_VVLVA_80_) and negatively charged residues preceding these (Fig 1A). To examine whether this SIM is a unique feature of S100A1, we performed an identical prediction procedure for other S100A family proteins (Table S3). We found that four of the sixteen S100 proteins, including S100A1, S100A8, S100A12 and S100A16, contain putative SIM sequences. Surprisingly, there was little overlap between our prediction and a previous SUMO:SIM interaction screen [35]. This discrepancy could be due to the different sensitivities of the prediction algorithms, as lowering the stringency parameters in the GPS-SUMO 2.0 tool predicted several more S100 proteins to contain SIM sites (data not shown).

**Figure 1:**
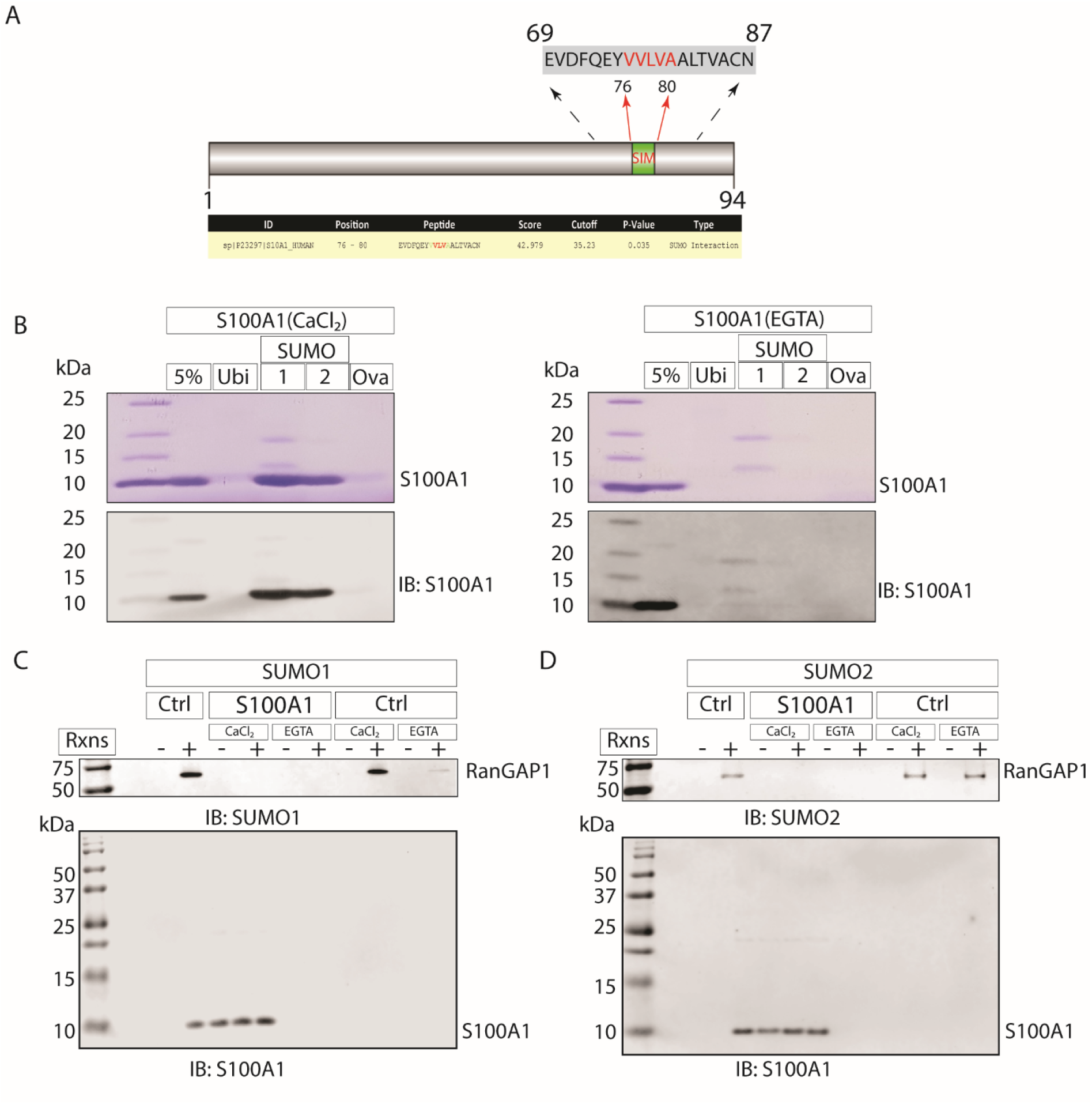
S100A1 binds SUMO but is not SUMOylated. A) Schematic representation of the S100A1 sequence showing the SIM (SUMO-interaction motif) predicted by GPS-SUMO in the C-terminal with the core of the SIM highlighted in red. B) S100A1 binds SUMO proteins in the presence of 1 mM Ca2+. SDS-PAGE (above) and Western blotting (below) analysis of a S100A1/SUMO pulldown assay performed in the presence of CaCl2 (left) or EGTA (right). Ubiquitin (Ubi) and ovalbumin (Ova) are included as controls. 5% represents the amount of S100A1 reserved as input from the total 10 µg S100A1 used in the assay. C-D) Western blotting analysis of in vitro SUMOylation assay (C: SUMO1; D: SUMO2) conducted at 37°C for 1 hr in the indicated conditions using antibodies directed against SUMO1, SUMO2 and S100A1. The assay control, RanGAP1, is sumoylated independently of modifying the assay protocol by addition of 2 mM CaCl2 or 2 mM EGTA in the assay buffer.

In order to experimentally validate the *in silico* prediction and determine whether S100A1 can interact with SUMO proteins, we performed an *in vitro* SUMO binding assay (SUMO interaction assay) using recombinantly purified S100A1 and SUMO1 or SUMO2 immobilized on sepharose beads in the presence of either Ca^2+^ or EGTA. The SUMO interaction assay showed that S100A1 specifically interacts with SUMO1 or SUMO2 in the presence of Ca^2+^ (Fig 1B, left), but not with the control proteins ubiquitin or ovalbumin. In contrast, the presence of EGTA completely prevented the interaction of the SUMO proteins with S100A1 (Fig 1B, right).

We also assayed whether S100A1 can be SUMOylated using an *in vitro* SUMOylation assay, again employing recombinantly purified S100A1. RanGAP1, a classical SUMO1 target [66, 67], served as a positive control and demonstrated strong SUMOylation with SUMO1 in the presence of Ca^2+^, which was weakened with EGTA (Fig 1C, D, Fig S1). However, using original assay conditions (without Ca^2+^ or EGTA) demonstrated a strong SUMOylation of RanGAP1 (Fig 1B, C, Fig S1). For SUMOylation of RanGAP1 with SUMO2, addition of either Ca^2+^ or EGTA had little impact. In contrast, SUMOylation of S100A1 did not occur in the presence or absence of either Ca^2+^ or EGTA.

Thus, the experiments corroborated the GPS-SUMO 2.0 prediction and further showed that the interaction of S100A1 with SUMO1 and SUMO2 is Ca^2+^ dependent. In contrast, the SUMOylation assay demonstrated that S100A1 cannot be SUMOylated *in vitro* by either SUMO1 or SUMO2.

### SUMO1 increases S100A1 protein abundance

Although less explored than SUMOylation, SUMO protein:protein interaction has been demonstrated to regulate a variety of biological processes [35, 68]. To assess the functional consequences of the SUMO:S100A1 interaction, we overexpressed MYC-tagged or non-tagged S100A1 together with FLAG-tagged SUMO1 in neonatal rat ventricular cardiomyocytes (NRVM) as well as in COS1 cells (Fig 2A, Fig S2A). Western blot analysis revealed that S100A1 and SUMO1 co-overexpression led to a significant increase in S100A1 protein levels (Fig 2B). In contrast, S100A1 co-overexpression with SUMO1 did not change the mRNA expression of S100A1 compared to co-expressed GFP (Fig 2C).

**Figure 2:**
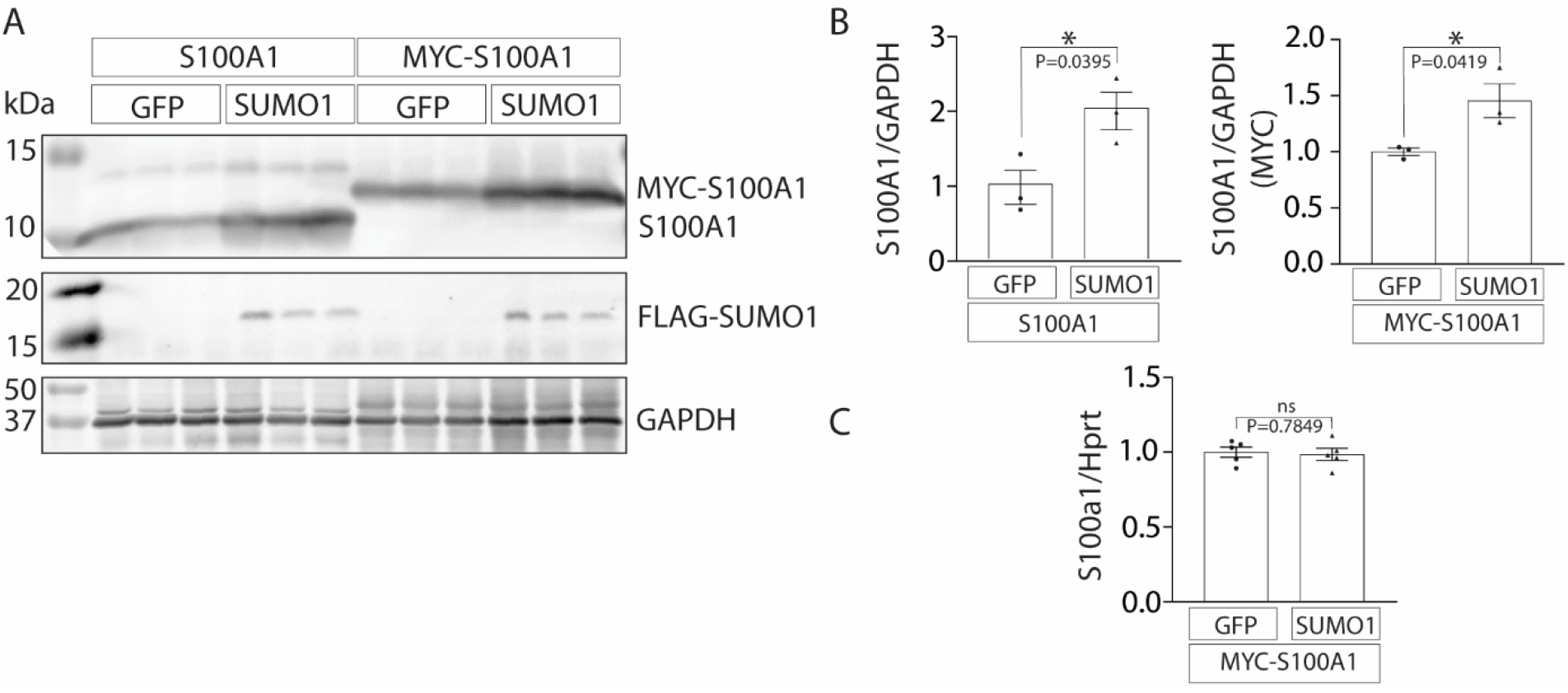
S100A1 and SUMO1 co-(over) expression increases S100A1 protein abundance in NRCM. A) untagged (left) or MYC-tagged (right) S100A1 was co-overexpressed together with either GFP or SUMO1 in NRVM using adenoviral gene delivery. 24 hrs post NRVM adenoviral transduction, cells were processed for western blotting analysis and probed with antibody directed against S100A1, FLAG and GAPDH. B) Quantification of S100A1 protein fold change normalized to GAPDH (left: untagged; right: MYC-tagged) co- (over) expressed together with either GFP or SUMO1. Values are presented as mean ± SEM., n=3 (independent samples). C) Fold change in levels of mRNA for S100A1 in cells co-overexpressing S100A1 together with either GFP or SUMO1. Values are presented as mean ± SEM., n=5 independent samples. Statistical analysis was done using unpaired two-tailed *t*-test; P<0.05 was considered significant. ns=not significant

Together, these results suggest that SUMO1 regulates S100A1 protein expression at the post-transcriptional level.

### The SIM of S100A1 is essential for S100A1 post-transcriptional stability

Most non-covalent SUMO:target protein interactions occur through the conserved SIM [30, 69]. To test if the SIM in S100A1 is required for SUMO1:S100A1 interaction and to regulate S100A1 protein expression, we generated N-terminally MYC-tagged S100A1 truncation mutants that lacked the predicted SIM (S100A1-1-74 containing residues 69-74 and lacking the SIM, Fig 3A).

**Figure 3:**
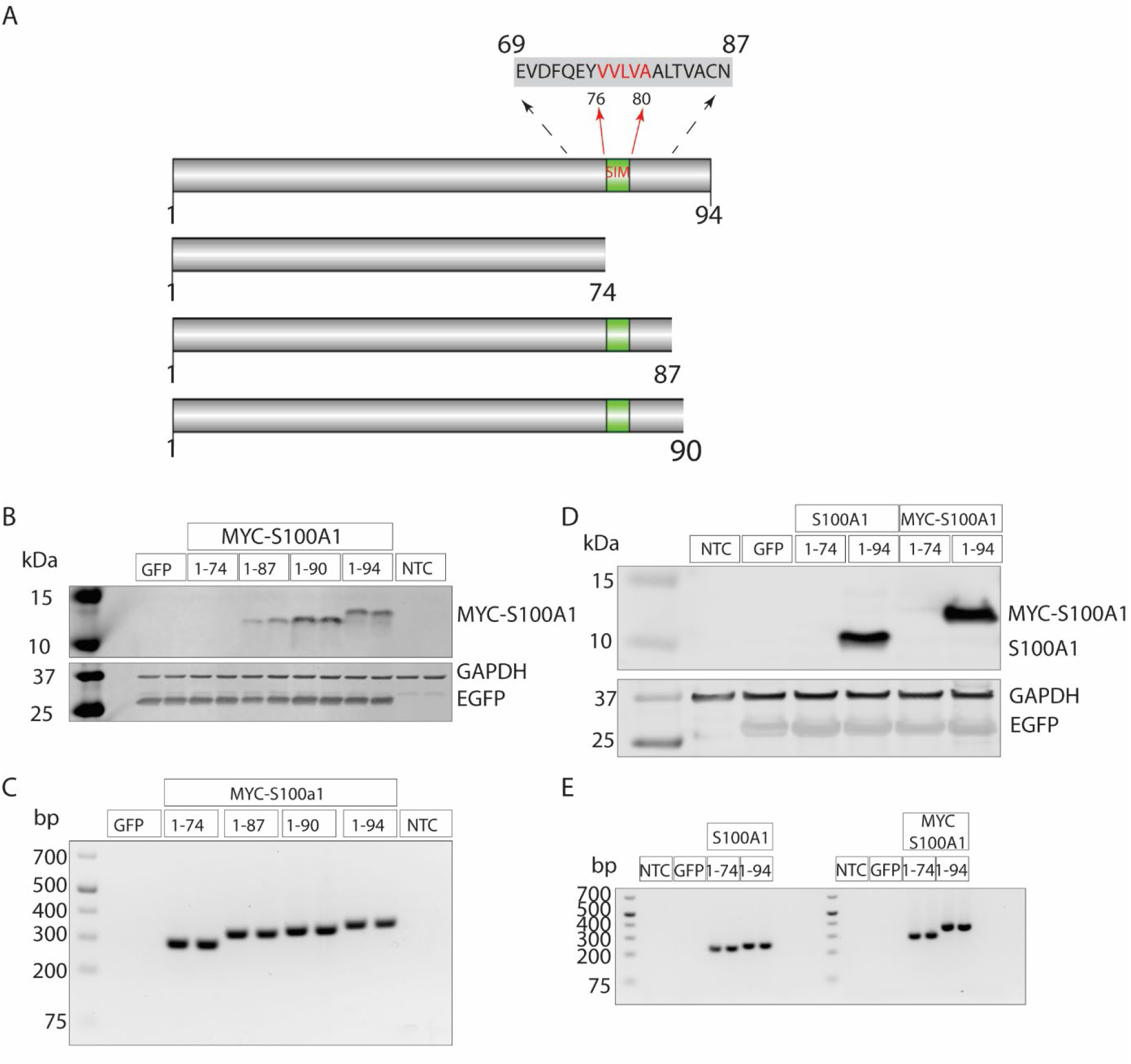
Truncated S100A1 protein lacking the SIM is unstable. A) Schematic representation of S100A1 and S100A1 truncation mutants designed based on the GPS-SUMO prediction of the SIM (SUMO-interaction motif). S100A1-1-94, S100A1 truncation mutants; S100A1-1-74 (containing residues 69-74 and lacking the SIM), S100A1-1-87 (containing residues 69-87) and S100A1-1-90 (containing residues 69-87 plus 3 aa). B) Western blotting analysis of COS1 cells transfected with MYC-tagged S100A1 and truncation mutants; EGFP and NTC (non-transfected control cells). 48 hrs post transfection, samples were processed for western blotting and probed with antibodies directed against MYC, GAPDH and GFP. The S100A1-1-74 mutant could not be detected at the protein level. C) End-point PCR analysis using the same experimental setup as in B shows that all the constructs could be detected at the mRNA level. D) Western blotting analysis of NRVM cells transduced with either untagged or MYC-tagged S100A1 (1-74) and S100A1(1-94); and the control EGFP as well as NTC. 24 hrs post transduction, samples were processed for western blotting and probed with antibodies directed against S100A1, GAPDH and GFP. The S100A1-1-74 mutant could not be detected at the protein level, independent of the presence or absence of the MYC-tag. E) The same experiment as in D was repeated and samples were processed for end-point PCR to detect whether the transgene is expressed at the mRNA level. All the constructs could be detected at the mRNA level.

Surprisingly, after overexpression in COS1 cells, S100A1-1-74 could not be detected on the protein level (Fig 3B). We then generated additional truncation mutants that contained residues 69-87 (S100A1-1-87) or an additional 3 amino acids (S100A1-1-90). S100A1 protein expression was slightly decreased in the S100A1-1-87 mutant, and indistinguishable from full length S100A1 (S100A1-I-94) in the S100A1-1-90 mutant (Fig 3B). In contrast, all truncation mutants were detected at the mRNA level (Fig 3C) and all expression plasmids used demonstrated similar GFP expression (Fig S3B).

To test if the MYC-tag affects protein stability of S100A1 mutant lacking the SIM, we then overexpressed S100A1-1-74 and S100A1-1-94 with and without the MYC-tag in NRVM. As for the experiments in COS1 cells, S100A1-1-74 could only be detected at the mRNA level, whereas S100A1-1-94 could be detected both at the protein level and the mRNA level, regardless of the presence or absence of the MYC-tag (Fig 3D, E). Of note, transduction efficiency did not differ among transgenes (Fig S3C).

The core hydrophobic residues of the SIM (corresponding to residues 77-79 of S100A1) have been shown to directly interact with SUMO [32, 70, 71]. To corroborate results from S100A1-truncation mutant experiments, we next generated N-terminal MYC-tagged S100A1-SIM mutants either by deletion or amino acid substitution of the core residues of the SIM. The SIM deletion mutant (DSIM) lacks residues _77_VLV_79_ of S100A1, and the triple alanine (AAA) mutant was generated by replacing residues _77_VLV_79_ of S100A1 with three alanines (Fig 4A, residues 77-79 highlighted in red). These mutants were then overexpressed in COS1 cells. Similar to the SIM truncation mutant, the DSIM mutant could not be detected at the protein level (Fig 4B), whereas the AAA mutation led to a massive reduction in S100A1 protein levels (Fig 4B-C). In contrast, all mutants were equally expressed on the mRNA level (Fig 4D).

**Figure 4:**
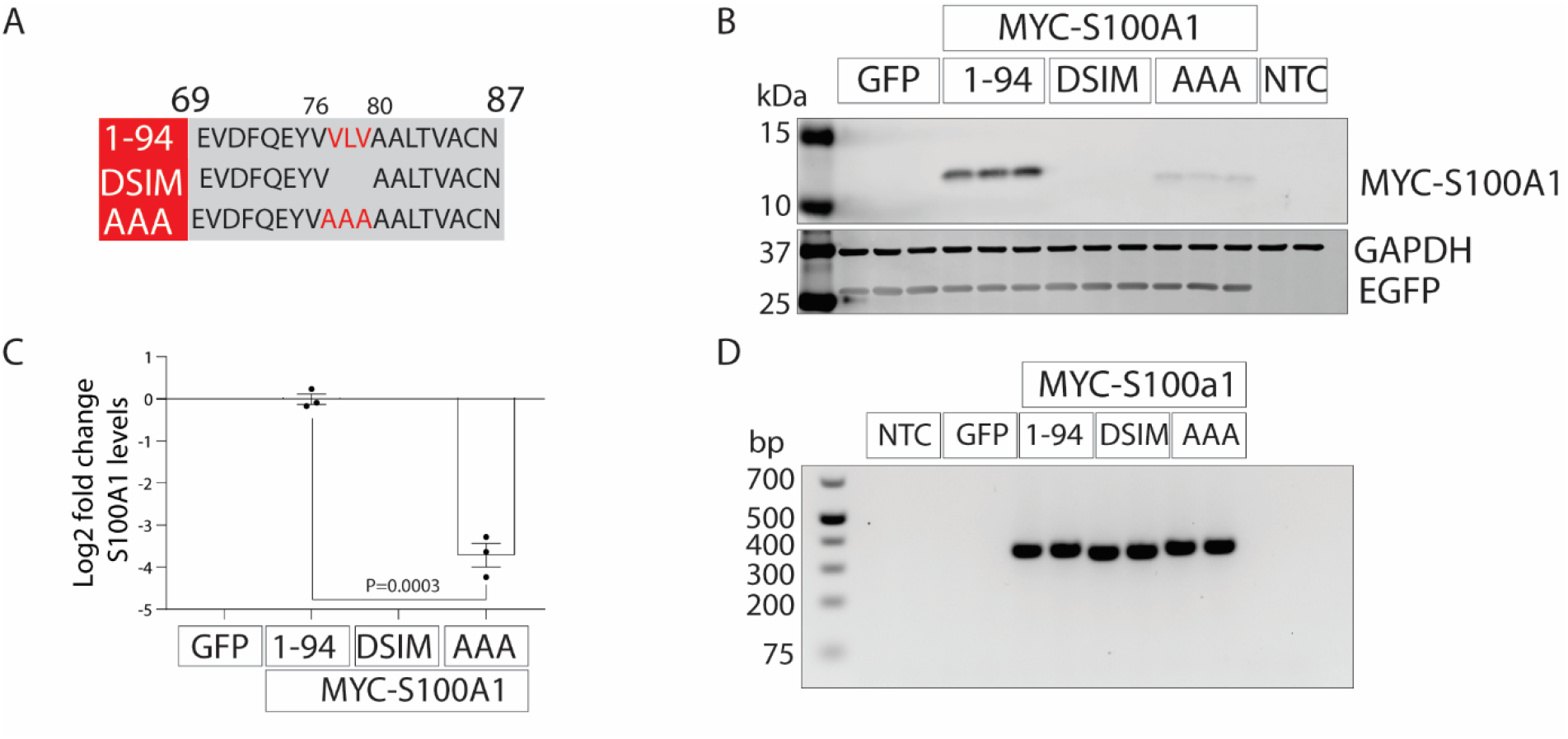
The SIM in S100A1 is critical for S100A1 protein stability. A) SIM sequence in S100A1 and the site directed modifications of the SIM core made by either deleting-_77_VLV_79_ (DSIM) or replacing the _77_VLV_79_ residues by three alanines (AAA) residues (at positions 77-79 in S100A1). B) Western blotting analysis of COS1 cells transfected with EGFP; MYC-tagged S100A1, DSIM and AAA; as well as NTC (non-transfected control cells). 48 hrs post transfection, samples were processed for western blotting and probed with antibodies directed against MYC, GAPDH and GFP. Western blotting analysis revealed that the DSIM mutant could not be detected at the protein level and the AAA mutant had dramatically reduced S100A1 expression at the protein level. C) Quantification of the S100A1 protein (1-94 and AAA mutant) normalized to GAPDH. The data are presented as a Log_2_ fold change with respect to S100A1 (1-94). Values are presented as mean ± SEM, n=3 independent samples. Statistical analysis was done using unpaired two-tailed *t*-test; P < 0.05 was considered significant. D) The same experiment as in B was repeated and samples were processed for end-point PCR to detect whether the transgene is expressed at the mRNA level. All the constructs could be detected at the mRNA level.

Together, these experiments suggest that the SIM in S100A1 is essential for S100A1 post-transcriptional protein stability.

### MG-132 treatment blocks degradation of the S100A1-SIM mutants

The lack of detectable S100A1 protein expression due to genetic SIM deletion or inactivation led us to hypothesize that these S100A1 variants are prone to immediate post-translational proteasomal degradation. To test this, we expressed the MYC-tagged S100A1 mutants in the presence or absence of MG-132. Western blot analysis revealed that proteasomal inhibition resulted in the stabilization of S100A1-1-74 and S100A1-DSIM proteins, and further increased the amounts of S100A1-1-94 and S100A1-AAA proteins (Fig 5A). Proteasomal inhibition also increased the expression level of GFP, while equal loading was verified by consistent endogenous GAPDH expression (Fig 5A).

**Figure 5:**
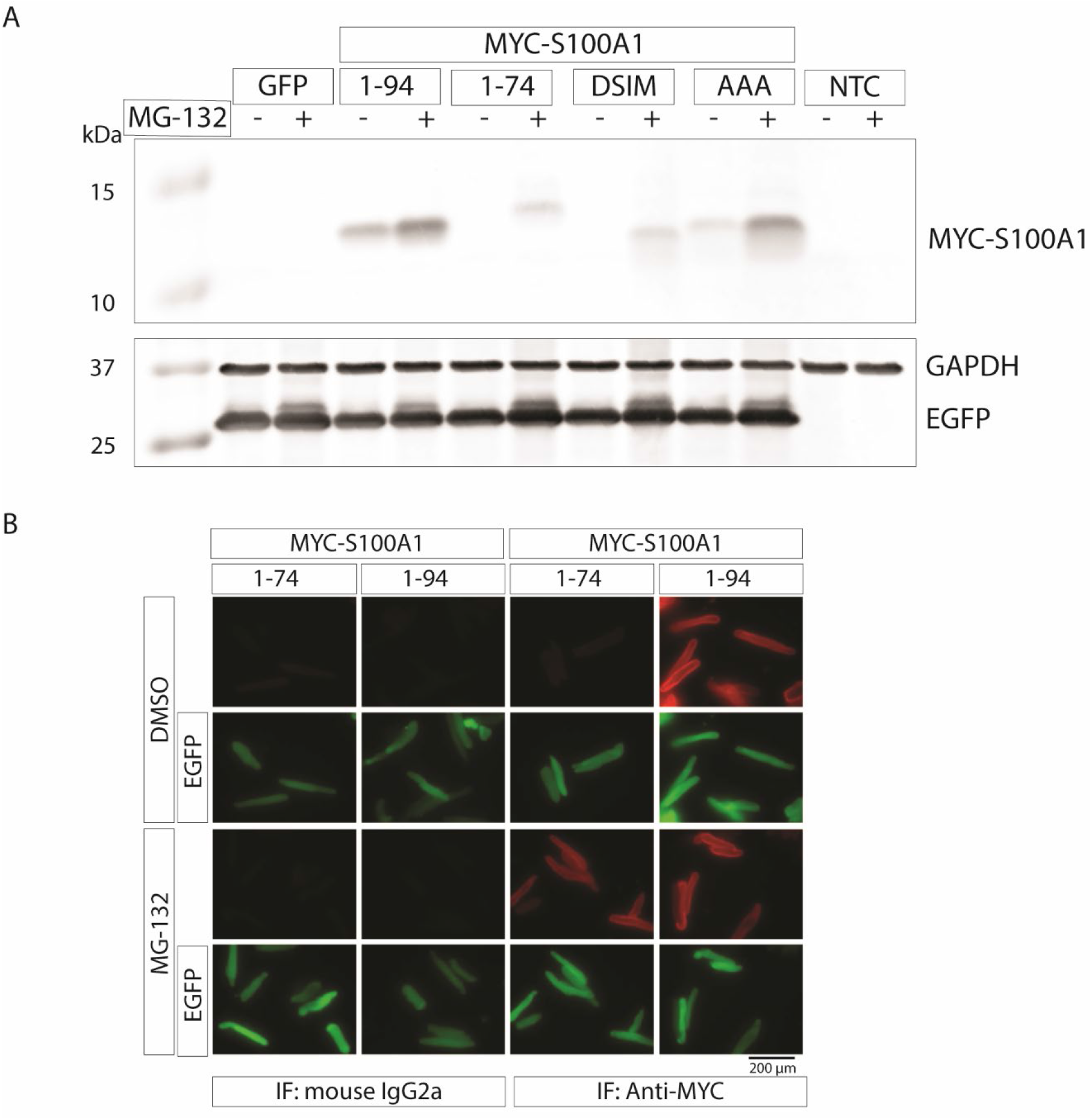
Proteasome inhibition blocked degradation of the S100A1 truncation mutant S100A1-1-74. A) Western blotting analysis of COS1 cells transduced with untagged S100A1-1-74 and S100A1-1-94 and the control EGFP in the absence or presence of 10 µM MG-132 (a proteasome inhibitor). 24 hrs post transduction, samples were processed for western blotting and probed with antibodies directed against S100A1, GAPDH and GFP. Western blotting analysis revealed that S100A1-1-74 could be detected at the protein level in the presence of 10 µM MG-132. B) For immune fluorescence (IF) analysis, ARVM (Adult rat ventricular myocyte) were transduced with MYC-tagged S100A1-1-74 and S100A1-1-94 in the absence or presence of 10 µM MG-132. ARVM were treated with either DMSO or 10 µM MG-132 for 10 hrs prior to fixation in 4 % PFA for the IF assay. The IF assay employed mouse anti-MYC antibody and as an isotype control mouse IgG2a. The IF assay revealed that MYC-tagged S100A1-1-74 could be detected at the protein level in the presence of 10 µM MG-132. Shown are representative images of anti-MYC staining (red) and EGFP fluorescence (green). Scale bar: 200 µm.

To exclude that the degradation of S100A1 devoid of the SIM is a post-harvest artifact and due to sample processing in the lysis buffer, we adenovirally overexpressed MYC-tagged S100A1-1-74 and S100A1-1-94 in adult rat ventricular myocytes (ARVM) in the presence or absence of MG-132 and performed immunofluorescence staining. Similar to the western blot results, S100A1-1-74 could only be detected after proteasomal inhibition by MG-132. In contrast, S100A1-1-94 could be detected independently of the presence of MG-132 (Fig 5B).

In summary, our experimental data indicate that S100A1 can non-covalently interact with SUMO1/2 and that the SIM of S100A1 is critical for S100A1’s post-transcriptional stability and to oppose proteasomal degradation. To advance our understanding of the function-structure relationship, we next sought to further explore the S100A1:SUMO interaction mode.

### Computational molecular docking suggests a novel SIM interaction mode in the S100A1:SUMO1 complex

To predict the structural mode of the SUMO1:S100A1 interaction, we applied HADDOCK, an information-driven, flexible protein-protein docking tool [43]. This molecular docking procedure was driven by ambiguous interaction restraints that encoded the results obtained with GPS-SUMO2.0 and the mutagenesis experiments.

We first docked SUMO1 to the WT S100A1 protein. Based on the results of the SUMO interaction assays, we selected the Ca^2+^-bound S100A1 structure for docking. We assumed that S100A1 is mainly present as a homodimer, which is supported by evidence from previous biochemical and structural studies on S100A1 and homologues [46, 47, 72-76]. For docking, we defined residues _76_VVLVA_80_, located in the SIM in H-IV of each subunit of the S100A1 homodimer, as active residues, as they were predicted to make direct interactions with SUMO and because mutation of residues 77-79 adversely affected the detected protein level. Interestingly, the helical parts of the two S100A1 monomers that contain the SIMs are situated directly adjacent to each other in an antiparallel arrangement in the homodimeric structure, making a hydrophobic base at the center of an “arched cleft”, formed by parts of the canonical C-terminal EF-hand motifs (Fig 6A).

**Figure 6:**
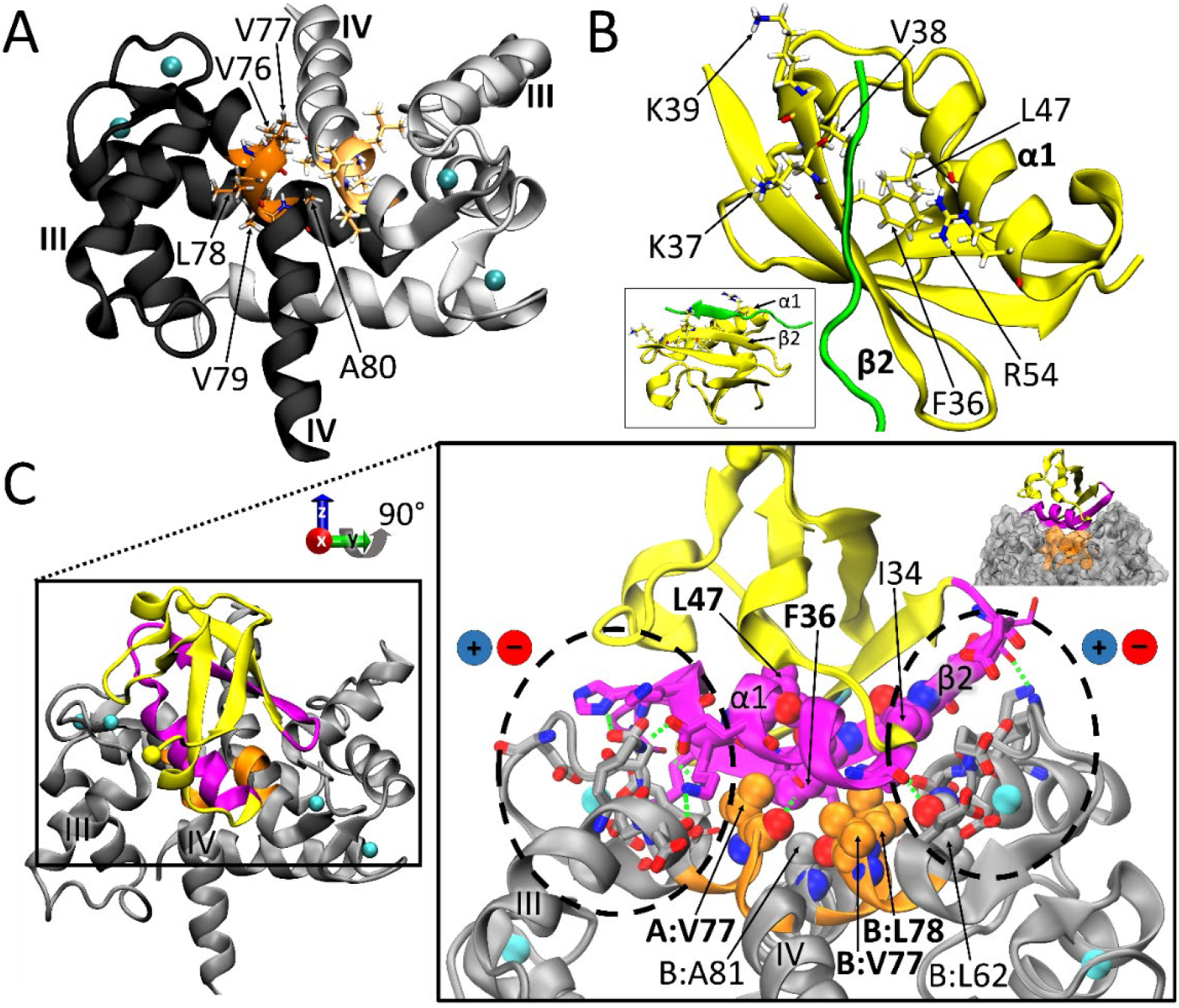
Structures of the proteins used for docking and predicted docked complex of S100A1 and SUMO1. A) Holo-S100A1 homodimer (PDB id, NMR model 1) shown in cartoon representation. Calcium ions are shown as cyan spheres. The first S100A1 subunit (chain A) is shown in dark grey, the second subunit (chain B) in light grey. Helices III and IV are indicated by labels. The SIMs are shown in dark orange (chain A) and light orange (chain B) in the antiparallel helices IV, and SIM residues are depicted as sticks colored by atom-type and labeled in chain A. B) SUMO1 (PDB id 6V7Q, crystal structure) in yellow cartoon representation. The bound peptide ligand, present in the structure, is shown in green cartoon representation. The α1 helix and β2 strand are indicated by labels. The apolar and positively charged sidechains mentioned in the text are shown in stick representation and labeled accordingly. The small inlay in the bottom left shows a side view of the respective peptide-SUMO1 complex. C) Complex of SUMO1 and S100A1 predicted by docking with HADDOCK. The best-scoring pose of the best HADDOCK cluster is displayed in two approximately perpendicular views. The S100A1 homodimer is shown in grey cartoon representation and the two SIMs are highlighted in orange. SUMO1 residues comprising the set I active residues are shown in magenta, the remaining SUMO1 residues are shown in yellow cartoon representation. The N- and C-termini of the SUMO1 structure (residues 20-94) are indicated by Cα atoms shown as yellow spheres. Ca^2+^ ions are shown as cyan spheres. Residues that form interactions (i.e. intermolecular contacts provided by Prodigy or polar contacts computed by PyMOL) at the SUMO1-S100A1 dimer interface within 5.5 Å are shown as either sticks (involved in potential salt-bridges and hydrogen bonds) or spheres (involved in potential apolar contacts; additionally indicated by labels, with apolar residues also shown in panel A or B as well highlighted in boldface) (colored by atom-type), with hydrogen bonds highlighted by green dashed lines. Dashed circles emphasize the clustering of potential polar contacts and charged interactions between the S100A1 dimer and SUMO1 within the predicted binding mode as discussed in the text. An additional alternative view of the predicted binding mode highlighting the surface of the S100A1 dimer is shown in the upper right inset.

For SUMO1, we used two alternative sets of active residues, set I and II (see ‘Experimental procedures’ for the exact residue definitions). Set I included residues in the hydrophobic groove formed by the α1-helix and the β2-strand of SUMO1 (plus the loop connecting them), which represents the canonical binding site that embeds classical SIMs, normally binding to the hydrophobic groove in SUMO in an extended β-strand-like conformation (Fig 6B) [30]. These residues include the crucial hydrophobic residues F36, V38, L47 at the bottom of the groove, and the positively charged residues K37, K39, R54 at the side, which are thought to play a role in electrostatic interactions with the negatively charged residues of the interaction partners containing a SIM, such as SUMO-interacting peptides [68]. Because the structure of the S100A1 dimer contains a SIM in a helical rather than an extended structure (which would be expected for a classical SIM), we also tested an alternative set of active residues for SUMO1, termed set II. Set II is a combination of SUMO1 residues that have been termed Class II SUMO interactions and have been found to be involved in noncovalent SUMO1-protein interactions (SUMO1-Ubc9 and SUMO1-DPP9) [77]. They differ from the canonical Class I binding via a SIM, being located on the opposite side of the groove formed by the SUMO1 α1-helix and β2-strand [78]. We observed that the conformational variability amongst the 20 NMR structure models (PDB id 2LP3) in the relative orientation of the two canonical C-terminal EF-hand motifs that form the boundaries of the “arched cleft” in the S100A1 dimer affects the accessibility of the SIMs (Fig S5). We therefore conducted separate dockings to all the representatives obtained from conformational clustering of the NMR models (labeled holo/apo-NMR cluster 0-4, accordingly). Comparing the docking results with the active residues from set I and set II shows that the HADDOCK scores are always better for the dockings employing set I, which corresponds to SUMO1 residues comprising the SIM-binding groove, even when restraint violation energies are subtracted from the final scores (see Table S4 vs. Table S5). Based on the best docking prediction with set I (‘WT, holo-NMR cluster 2’, Table S4), SUMO1 associates with the S100A1 dimer by covering parts of the top of the “arched cleft” that are lined by residues from both SIMs of the S100A1 dimer with its hydrophobic cleft, thus exhibiting a tilted contact with the S100A1 dimer (Fig 6C). In this binding mode, the hydrophobic cleft of SUMO1 covers one SIM almost completely, while the second neighboring SIM is partially covered. This binding mode therefore seems to maximize apolar contacts between the hydrophobic S100A1 SIM residues (A:V77, B:V77, B:L78, B:A81) and apolar residues in the hydrophobic cleft of SUMO1 (I34, F36, L47) (Fig 6C). Furthermore, the docking indicates that SUMO1 can interact with S100A1 via polar and charged interactions in this binding mode, mainly between residues in the C-terminal EF-hands of both S100A1 subunits and residues in the α1-helix and the β2-strand (plus the corresponding connecting loop) of SUMO1, with the EF-hands thus “grasping” SUMO1 in a “pincer-like” mode (Fig 6C). Also, the N- and C-terminal ends of SUMO1, which were not considered during docking due to their flexibility, have enough space to protrude into the solvent and not adversely interfere with the binding mode (Fig 6C). Interestingly, the S100A1 dimer conformation in this pose (‘holo-NMR cluster 2’, which corresponds to model 9 of 2LP3) shows the greatest accessibility of the SIM amongst the NMR conformational clusters (Fig S5). Furthermore, docking of SUMO1 to this S100A1 dimer conformation yields only two docking clusters with similar HADDOCK scores, corresponding to two complexes that are symmetric with respect to SUMO1 binding, which is consistent with the symmetric structure of the S100A1 homodimer (Table S4 and Fig S6).

For comparison, the best-ranking docking modes obtained for set I and set II active residues are shown in Fig S7. Comparing the estimated binding affinities obtained from re-scoring the dockings with set I and set II active residues by applying FoldX5.0 and Prodigy shows that the predicted pose for the set II active residues have a much weaker affinity than the poses generated with set I (Table 1; first row vs. second row). The estimated binding affinities therefore corroborate the interaction of the SIM with the α1-helix and the β2-strand of SUMO1 (set I) being favored over the interaction with residues in loops on the opposite side of the SUMO1 protein (set II).

**Table 1:**
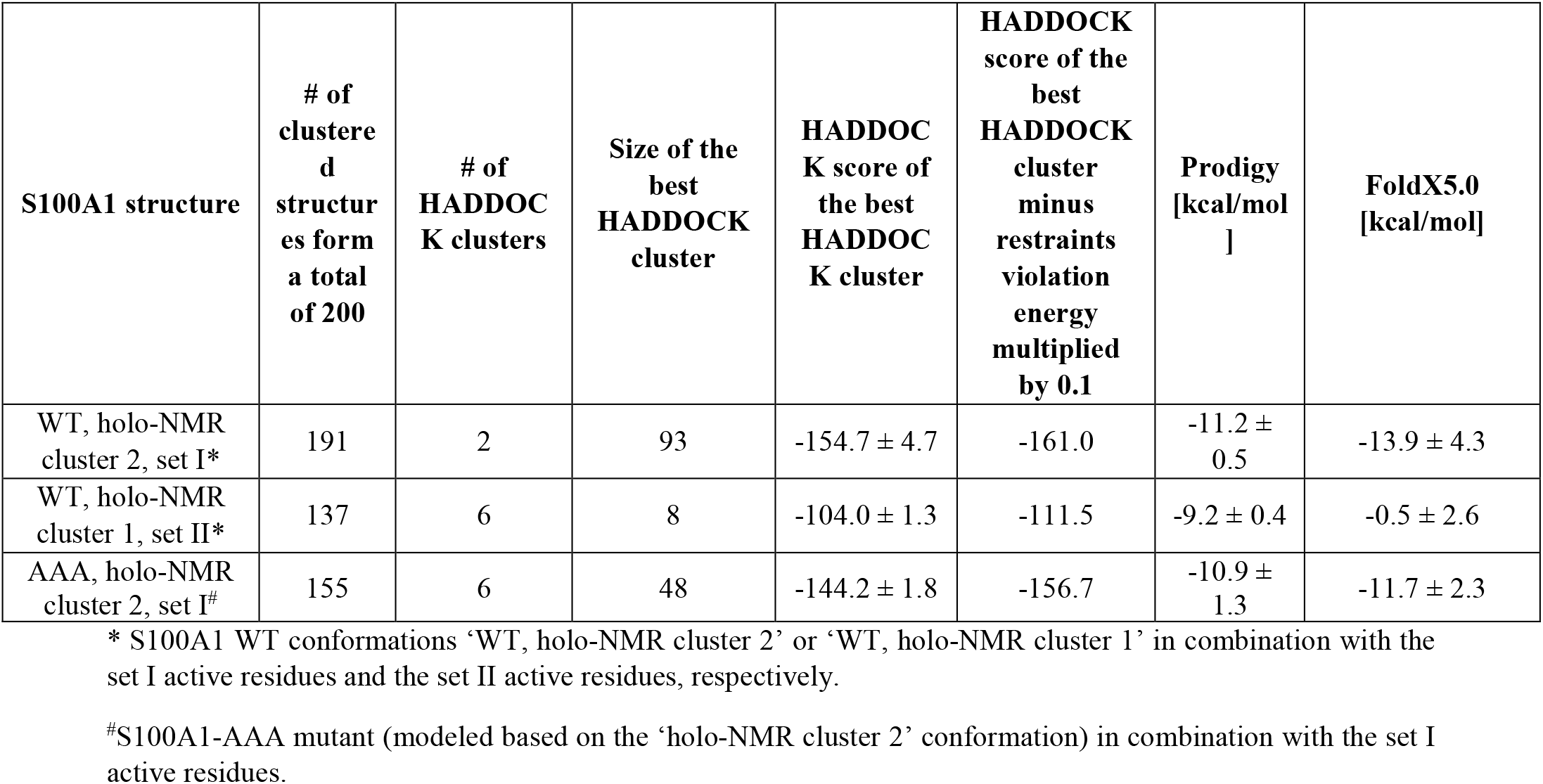
Summary of HADDOCK docking performance parameters for docking SUMO1 to the S100A1 homodimer. Binding affinity estimates from Prodigy and FoldX5.0 are given as averages over the four top-scoring poses of the best HADDOCK cluster (with the HADDOCK score employed to rank the HADDOCK clusters).

| **S100A1 structure** | **# of clustered structures from a total of 200** | **# of HADDOCK clusters** | **Size of the best HADDOCK cluster** | **HADDOCK score of the best HADDOCK cluster** | **HADDOCK score of the best HADDOCK cluster minus restraints violation energy multiplied by 0.1** | **Prodigy [kcal/mol]** | **FoldX5.0 [kcal/mol]** |
| --- | --- | --- | --- | --- | --- | --- | --- |
| WT, holo-NMR cluster 2, set I* | 191 | 2 | 93 | -154.7 ± 4.7 | -161.0 | -11.2 ± 0.5 | -13.9 ± 4.3 |
| WT, holo-NMR cluster 1, set II* | 137 | 6 | 8 | -104.0 ± 1.3 | -111.5 | -9.2 ± 0.4 | -0.5 ± 2.6 |
| AAA, holo-NMR cluster 2, set I<sup>#</sup> | 155 | 6 | 48 | -144.2 ± 1.8 | -156.7 | -10.9 ± 1.3 | -11.7 ± 2.3 |

\* S100A1 WT conformations ‘WT, holo-NMR cluster 2’ or ‘WT, holo-NMR cluster 1’ in combination with the set I active residues and the set II active residues, respectively.
<sup>#</sup>S100A1-AAA mutant (modeled based on the ‘holo-NMR cluster 2’ conformation) in combination with the set I active residues.

We next docked SUMO1 to the apo-S100A1 dimer structure (i.e., with no Ca^2+^ bound; further referred to as apo-dockings), using the same preparation procedure and settings as for holo-S100A1, by applying the set I active residues. A similar best-scoring pose was obtained, see Table S6 and Fig S8. Binding affinity estimates comparing SUMO1 dockings to the apo- and holo-S100A1 dimer structure indicate more favorable binding to holo-S100A1 compared to apo-S100A1 (see Table S7). Comparison of the apo- and the holo-S100A1 dimer structures shows that the SIMs are much more exposed in the holo-than in the apo-S100A1 dimer structures (Fig S8C), likely facilitating binding of SUMO1 to a holo-S100A1 conformation. These differences provide a possible reason for the Ca^2+^-dependent interaction of SUMO1 and S100A1 found in the SUMO interaction assays[46, 47, 73-76].

The complementarity between the electrostatic potentials of the holo-S100A1 dimer (‘holo-NMR cluster 2’) and SUMO1 at the predicted docking interface (see Fig S9) further corroborate the HADDOCK models. Strikingly, *de novo* prediction of diffusional encounter complexes between the holo-S100A1 dimer (‘holo-NMR cluster 2’) and SUMO1 by applying Brownian dynamics-based docking using SDA (Simulation of Diffusional Association) identified two diffusional encounter complexes (SDA-cluster 5 and 6; see Table S8) within the top-4 docking modes (as regards SDA-cluster size and average interaction energy) that are similar to the two symmetric HADDOCK binding modes (see Fig S10). Thus, the SDA results further support our HADDOCK models.

In conclusion, the S100A1:SUMO1 docking mode obtained suggests a novel SUMO1-SIM interaction mode.

### The SIM in S100A1 is predicted to facilitate SUMO:S100A1 binding

To explore the effect of the AAA triple mutation on the S100A1:SUMO1 interaction, we also docked SUMO1 to the S100A1-AAA mutant (modeled based on model 9 from the S100A1 structure 2LP3, i.e., ‘holo-NMR cluster 2’) using HADDOCK. As this S100A1 mutant lacks a functional SIM, we hypothesized that it would be unable to interact with SUMO1.

As alanine is known to be a helix-stabilizing residue, the alanine mutations were not expected to significantly alter the S100A1 conformation upon mutation. The predicted binding modes of SUMO1 to the S100A1-AAA dimer are similar to those for the WT S100A1 dimer (Fig S7C) but the score for the top-ranked cluster was less favorable (Table S1). As the HADDOCK score is designed to rank different poses of the same system rather than to discriminate binding affinities between different systems, we computed binding affinity approximations with FoldX5.0 and Prodigy. We could observe a trend in the estimated binding affinities, indicating that the S100A1-AAA homodimer interacts less strongly with SUMO1 than the WT S100A1 dimer (Table 1).

## Discussion

In the present study, we demonstrate a novel regulatory mechanism that controls S100A1’s post-translational stability. We reveal a S100A1:SUMO1 interaction that stabilizes the S100A1 protein and find that the SIM in S100A1 is indispensable for preventing proteasomal degradation and is predicted to interact with SUMO via a novel structural binding mode. As such, this study advances our general understanding of the functional consequences of SUMO binding to SIM-containing proteins as well as our specific understanding of the regulation of S100A1 protein levels.

PTM by SUMO relies on the ability of SUMO to promote non-covalent protein:protein interactions, or the covalent conjugation of SUMO to target proteins [30]. In many cases, SUMO-binding proteins with SIMs are also covalent SUMO targets, which might indicate a functional connection between covalent and non-covalent SUMO binding targets [69, 79]. Guided by the bioinformatics prediction of a SIM in S100A1, we demonstrate that S100A1 interacts with SUMO1 and SUMO2 proteins, but is not subjected to covalent SUMOylation.

Interestingly, human SUMO1 and SUMO2 display variations in the residues that contact SIMs, having an impact on SUMO-SIM affinities [69]. In line with this, a recent study confirmed that, indeed, there is only a small fraction of SUMO interacting proteins that binds to SUMO irrespective of its isoform [35]. In contrast, we demonstrate that S100A1 can interact with both SUMO1 and SUMO2 in a Ca^2+^-dependent manner. The requirement of Ca^2+^ in the assay buffer for interaction with SUMO aligns with previous experimental findings that Ca^2+^ is required for the interaction of S100A1 with target proteins such as SERCA2a, RYR2 or ATP5A1 [21-23, 25, 26]. This Ca^2+^-dependency was further corroborated in the molecular docking analysis, where apo-S100A1 demonstrated a weaker binding affinity to SUMO1 than holo-S100A1.

We then focused on the consequences of SUMO1 and S100A1 interaction. Overexpression of SUMO1 together with S100A1 increased S100A1 protein, but not transcript levels, suggesting increased protein stability. Traditionally, regulation of protein stability has been linked to SUMOylation, but not to non-covalent SUMO interaction [80-82]. To our knowledge, this is the first report linking SUMO:SIM interaction to protein stability, emphasizing on the novelty of this finding.

The impact of the SIM in S100A1’s post-translational stability was assessed by employing truncation mutations and site directed mutagenesis of the core of the SIM. These mutant S100A1 proteins demonstrated accelerated degradation, although the respective transcript expression was not affected. This phenotype could be rescued by proteasomal inhibition, which stabilized protein levels. Thus, enhanced proteasomal degradation is most likely contributing to the decreased post-translational stability. This finding provides the first evidence that the core residues of the SIM of S100A1 are indispensable for post-translational stability of S100A1. Furthermore, our findings uncovered a new function of SUMO:target protein interaction, that might regulate protein stability and oppose proteasomal degradation. If this is specific for S100A1 or extends to other SUMO interacting proteins however needs further validation.

Integrative modeling of the S100A1:SUMO interaction revealed a potentially novel SIM interaction mode between SUMO1 and the two SIMs present in the S100A1 homodimer. Normally, a peptide stretch that contains a SIM is assumed to bind to the groove of SUMO1 in an extended conformation, mediated via weak backbone interactions, apolar and electrostatic interactions, as observed in experimentally determined structures of peptide:SUMO1 complexes in the Protein Data Bank (PDB) [83] (e.g., 2ASQ [71] and 2LAS [84], also see Fig S11) [30]. However, in the structures of the S100A1 homodimer (e.g., 2LP3 and 5K89 [73]), the predicted SIM is in H-IV. Moreover, previous predictions and molecular dynamics simulations for a peptide corresponding to the C-terminal H-IV helix of S100A1 (S100A1ct), which also contains the SIM, show that this region adopts a helical conformation in aqueous solution and do not support the idea that the SIM would adopt a significantly populated extended conformation [85]. Intriguingly, upon structural superposition of the SUMO1 proteins, A:V77 and B:L78 (in separate SIMs, but in close proximity in the homodimer) roughly align with the two apolar residues that mainly occupy the hydrophobic cleft of SUMO1 in the structures of peptide:SUMO1 complexes (2LAS and 2ASQ) (Fig S11), suggesting that the S100A1 homodimer could mimic parts of a classical SIM:SUMO interaction: apolar interactions between SIM residues and residues in the cleft between the α1-helix and the β2-strand of SUMO, as well as interactions between negatively charged SIM residues and SUMO. It is important to note that our findings are based on the assumptions that SIM residues are part of the interaction interface and that the folded native S100A1 homodimer structure is the target of SUMO1. The former assumption is supported by the mutation data and Brownian dynamics simulations of protein-protein diffusional encounters that did not include any interaction restraints. The latter assumption is supported by the observed Ca^2+^-dependence of SUMO1 binding. However, if the S100A1 monomer would be the target of SUMO, it could be (partially) unfolded, e.g., due to the exertion of some form of external force, or as a nascent S100A1 protein. Furthermore, the SIM overlaps with a region that has been identified to be important for dimerization in S100A4 [86]. So potentially lower dimerization propensities could at least partially contribute to lower interactions of SUMO with S100A1 mutants (as less S100A1 dimer would be present to interact with), and thus S100A1 stability.

One limitation of our study is that we could not experimentally test if the S100A1 SIM mutants can still interact with SUMO1 due to the instability of the SIM protein mutants. However, based on the molecular docking of S100A1-AAA with SUMO1, we find that the binding affinity is reduced compared to the WT S100A1:SUMO1 interaction. This suggests that the identified SIM in S100A1 is required for the S100A1:SUMO1 interaction. One possible mechanism through which the SIM-mediated S100A1:SUMO interaction could stabilize S100A1 levels is by limiting accessibility to ubiquitination [87]. How this exactly occurs is still unclear, and this mechanism warrants further investigation.

What is the biological function of the S100A1:SUMO1 interaction? SUMO1 is downregulated in human and experimental heart failure models [88]. Thus, based on our findings reported here, downregulation of SUMO1 could contribute to the reduced S100A1 protein expression that has been described in various heart failure models that drives progression of the disease and worsens survival [8-11]. This could provide an additional mechanistic layer that contributes to the dysregulation of S100A1 protein expression during chronic stresses. In this context, SUMO1 overexpression in isolated cardiomyocytes increased Ca^2+^ handling and rescued cardiac dysfunction in an experimental heart failure model [88]. These effects have been attributed to improved SERCA2a function, but it is tempting to speculate that increased SUMO1 levels could also stabilize S100A1 protein expression in the failing heart, which demonstrates overlapping effects *in vitro* and *in vivo* [9, 25, 89-92]. However, this SUMO1-S100A1 crosstalk in the failing heart needs further validation.

Furthermore, SUMO conjugation has been linked to promoting the assembly of protein complexes, serving as a molecular glue, that facilitates the interaction of functionally related proteins [68]. For this, the promotion of protein interaction by SUMO conjugation is typically achieved through SUMO-dependent recruitment of binding partners, which contain distinct SIMs [29, 93]. Interestingly, among S100A1 target proteins, SERCA2a was identified to be SUMOylated [88]. Thus, it is tempting to speculate that the S100A1:SUMO1 interaction could facilitate the S100A1:SUMO1:SERCA2a interaction and play a role in the regulation of SERCA2a activity. Importantly, our molecular docking analysis would support this hypothesis, as both the N- and C-termini of SUMO are freely accessible for SUMOylation reactions when docked to S100A1. Furthermore, SERCA2a SUMOylation is reduced in heart failure [88]. Thus, impaired SERCA2a SUMOylation could result in weaker S100A1:SERCA2a interactions and exacerbate the defective Ca^2+^ handling in failing cardiomyocytes. In order to confirm these new hypotheses, further docking studies for all three proteins and experimental validation would be required but are beyond the scope of the current study.

In conclusion, our data shed new light on the post-translational control of S100A1 protein stability. We identified non-covalent modification by SUMO1 as a novel molecular checkpoint to regulate protein stability, and provide a structural model indicating that this could be mediated via a new type of SIM:SUMO1 binding. These observations also provide a rationale for generating S100A1 protein variants that hinder degradation and could have enhanced stability in the failing heart.

## Supporting information

zip for Fig 6

## Statements and Declarations

### Author contributions

ZHJ designed the study. ZHJ, AS, JZ and RS carried out experiments. ZHJ and PM analyzed the experimental data. MG and RCW designed, performed and interpreted the molecular modeling. ZHJ, MG, JR wrote the manuscript with assistance from all authors. FM, MB, RCW and PM provided research support and conceptual advice. All authors approved the final version of the paper.

### Data Availability

Additional data supporting findings of this study can be found in the supplementary material and are available from the corresponding author upon reasonable request. Furthermore, the HADDOCK and SDA poses displayed in the figures of this manuscript are available in the SI (docking_models.zip).

### Ethics approval

All animal procedures and experiments were performed in accordance with the ethical standards laid down in the 1964 Declaration of Helsinki and its later amendments institutional, guidelines of the University of Heidelberg and received approval from local authorities.

## Abbreviations (Non-standard abbreviations and acronyms)

SUMO: Small Ubiquitin-like Modifier
SIM SUMO: interacting motif
Ub: Ubiquitin
NRVM: Neonatal rat ventricular myocyte
ARVM: Adult rat ventricular myocyte
EGFP: Enhanced green fluorescence protein
DSIM: S100A1-SIM deleted mutant
CMV: cytomegalovirus
MOI: Multiplicity of infection
CM: cardiomyocyte
HFrEF: heart failure with reduced ejection fraction
PTM: post-translational modifications

## Acknowledgement

We thank Jasmin Kroemer, Stephanie Simon and Jennifer Birkenstock for technical help; We thank the late Jeffrey Robbins for providing us the FLAG-SUMO1 adenovirus construct. We thank Frauke Melchior for the expert help with in vitro SUMO interaction assay.

