## Supplementary material for "SUMO interacting motif (SIM) of S100A1 is critical for S100A1 post-translational protein stability": zip for Fig 6: README

This is part of the Supporting Information of the manuscript Jebessa, Z.H. et al. "SUMO interacting motif (SIM) of S100A1 is critical for S100A1 post-translational protein stability"

Manuel Glaser
Molecular and Cellular Modeling Group
Heidelberg Institute for Theoretical Studies (HITS)
Heidelberg, Germany

Web: https://www.h-its.org/research/mcm/

==========================================================================================
Overview of the models shown in figures related to the manuscript and offered for download
==========================================================================================

--------------------------------------------------
HADDOCK models of the docked S100A1-SUMO1 complex:
--------------------------------------------------

Naming scheme (elements are separated by underscores): 
* The first 'c2'/'c1'/'c4' etc. indicate the cluster id from the clustering of the S100A1 dimer NMR models used for docking
* 'set1'/'set2' indicates the active residues used for SUMO1
* 'cluster1'/'cluster2' indicates the HADDOCK cluster
* '_1_' indicates that the best-ranking pose of the corresponding HADDOCK cluster is used
* 'WT': indicating wild-type S100A1
* 'VLV77-79AAA': indicating the S100A1-AAA mutant
* 'aln': indicates that models have been aligned to the PDB id 2LP3 holo-S100A1 dimer NMR model 1
* 'ABC': indicates that the S100A1 subunits are named A and B (i.e., chain id) while the SUMO1 chain is referred to as C in the models
* Note: calcium atom names/residue names were renamed from CA+2/CA2 (HADDOCK naming scheme) to CA/CA in the models offered for download
* Important: please be aware that the numbering of S100A1 residues discussed in the manuscript corresponds to Uniprot numbering, but is different in the models (residues of both S100A1 subunits, including calcium ions if present, are consecutively numbered, the numbering of the chain 'A' is also shifted by 1 with respect to Uniprot numbering)
* run.cns is an example file containing the HADDOCK run parameters

HADDOCK models of predicted bound WT holo-S100A1 dimer-SUMO1 complexes:

* 'c2_set1_cluster1_1_WT_aln_ABC.pdb' (shown in Fig 6C, Fig S6A, Fig S7A, Fig S10A); the respective HADDOCK cluster-related data can be found in Table 1 and Table S4 (row label 'WT, holo-NMR cluster 2'), a summary of interface contacts can also be found in the footnotes of Table S4)

* 'c2_set1_cluster2_1_WT_aln_ABC.pdb' (shown in SI Fig 6B, Fig S10B); the respective HADDOCK cluster-related data can be found in the footnotes of Table S4

* 'c1_set2_cluster1_1_WT_aln_ABC.pdb' (shown in Fig S7B); the respective HADDOCK cluster-related data can be found in Table 1 and Table S5 (row label 'WT, holo-NMR cluster 1')

HADDOCK model resulting from docking SUMO1 to the WT apo-S100A1 dimer complex:

* 'c4_set1_cluster1_1_apo_ABC.pdb' (shown in Fig S8B); the respective HADDOCK cluster-related data can be found in Table S6 (row label 'WT, apo-NMR cluster 4')
 
HADDOCK model resulting from docking SUMO1 to the S100A1-AAA mutant: 

* 'c2_set1_cluster1_1_VLV77-79AAA_aln.pdb' (shown in Fig S7C); the respective HADDOCK cluster-related data can be found in Table 1 (row label, 'AAA, holo-NMR cluster 2') 

------------------------------------------------------------------------------
SDA models of S100A1-SUMO1 diffusional encounter complexes (shown in Fig S10):
------------------------------------------------------------------------------

* The PDB files containing the SDA cluster representatives of SUMO1 are labeled 'cl1.pdb' to 'cl10.pdb', the chain id is 'A'
(the corresponding cluster sizes, interaction energies of SDA-cluster representatives, average interaction energies of the SDA-clusters etc. can be found in Table S8)

* The corresponding PQR file of the holo-S100A1 dimer is named '2lp3_model_9.pqr' (S100A1 chain ids are 'A' and 'B') (important: as already described above for the HADDOCK models, the same considerations regards S100A1 residue numbering apply here)

* While the HADDOCK models are fully docked, the SDA models indicate intermediate states on the binding funnel.

* apbs.in and sda.in are example input files containing the APBS and SDA parameters
