## Supplementary material for "SUMO interacting motif (SIM) of S100A1 is critical for S100A1 post-translational protein stability": Suppl Information

*Jebessa et al.*

Supplementary Table S1: S100A1 and S100A1 mutants in pAdTrack-CMV vector used in overexpression experiments

| Name | Tag | type |
|---|---|---|
| **Deletion mutants** | | |
| S100A1-1-94 | none | Full length |
| S100A1-1-94 | Myc | Full length |
| S100A1-1-74 | none | Deletion - SIM lacking |
| S100A1-1-74 | Myc | Deletion - SIM lacking |
| S100A1-1-87 | Myc | Deletion - SIM intact |
| S100A1-1-90 | Myc | Deletion - SIM intact + 3aa |
| **SIM mutants** | | |
| S100A1-WT | Myc | Full length |
| S100A1-DSIM | Myc | aa $_{77}$VLV$_{79}$ deleted |
| S100A1-AAA | Myc | aa $_{77}$VLV$_{79}$ replaced by $_{77}$AAA$_{79}$ |

Supplementary Table S2: List of Primers used for qPCR and End-point PCR analysis

| Name | Sequence | purpose |
|---|---|---|
| Myc-S100A1-univ-Fwd | CAGAAACTCATCTCTGAAGAGG | End point PCR Myc-tagged Forward primer |
| S100A1-univ-Fwd | GACCCTCATCAACGTGTTCC | End point PCR non-tagged forward primer |
| S100A1-All-Rvs | TAGATCCGGTGGATCGGATATC | End point PCR all reverse primer |
| rHprt1-Fwd | CCAGCGTCGTGATTAGTGAT | qPCR house-keeping (Hprt1) forward primer |
| rHprt1-Rvs | AGAGGGCCACAATGTGAT | qPCR house-keeping (Hprt1) reverse primer |
| rS100a1-Fwd | TGAGCAAGAAGGAGCTGAAA | qPCR S100a1 forward primer |
| r S100a1-Rvs | CACCAGCACAACAAACTCCT | qPCR S100a1 reverse primer |

Supplementary Table S3: GPS-SUMO2 prediction of SIM for human S100 family proteins with "high" prediction threshold.

| # | Uniport ID | S100A | SIM | SUMOylation prediction |
|---|---|---|---|---|
| 1 | P23297 | 1 | yes | no |
| 2 | P29034 | 2 | no | no |
| 3 | P33764 | 3 | no | no |
| 4 | P26447 | 4 | no | no |
| 5 | P33763 | 5 | no | no |
| 6 | P06703 | 6 | no | no |
| 7 | P31151 | 7 | no | no |
| 8 | Q86SG5 | 7a | no | no |
| 9 | P05109 | 8 | yes | yes |
| 10 | P06702 | 9 | no | no |
| 11 | P60903 | 10 | no | no |
| 12 | P31949 | 11 | no | no |
| 13 | P80511 | 12 | yes | no |
| 14 | Q9HCY8 | 13 | no | no |
| 15 | Q9HCY8 | 14 | no | no |
| 16 | Q96FQ6 | 16 | yes | no |

Supplementary Table S4: Summary of HADDOCK docking performance parameters obtained by docking SUMO1 to five cluster representatives of the holo-S100A1 dimer structure with the set I active residues.

| **S100A1 structure** | **# of clustered structures from a total of 200** | **# of HADDOCK clusters** | **Size of the best HADDOCK cluster** | **HADDOCK score of the best HADDOCK cluster*** | **HADDOCK score of the best HADDOCK cluster minus the restraints violation energy multiplied by 0.1 (given in parentheses)** |
|---|---|---|---|---|---|
| WT, holo-NMR cluster 0 (representing 15 models) | 108 | 15 | 8 | -105.3 ± 6.3 | -119.4 (14.1) |
| WT, holo-NMR cluster 1 (representing two models) | 126 | 16 | 6 | -115.5 ± 15.7 | -121.5 (6.0) |
| **WT, holo-NMR cluster 2 (**representing one model)[§] | **191** | **2** | **93** | **-154.7 ± 4.7**[#] | **-161.0 (6.3)** [#] |
| WT, holo-NMR cluster 3 (representing one model) | 117 | 12 | 8 | -119.4 ± 10.2 | -145.8 (26.4) |
| WT, holo-NMR cluster 4 (representing one model) | 123 | 12 | 12 | -134.4 ± 6.7 | -145.7 (11.3) |

*The more negative the HADDOCK score is, the better the score is. The best scoring HADDOCK cluster is indicated by bold face. HADDOCK cluster scores are given as averages over the top-scoring four poses in the cluster, which is the default scoring approach.

[#]The HADDOCK score of the second-best HADDOCK cluster (size: 98) was -151.2 ± 1.6, and the HADDOCK score minus restraints violation energy was -157.7. This cluster had an approximately symmetric arrangement compared to the best HADDOCK cluster, corresponding to the two-fold symmetry of the S100A1 homodimer and is shown in Fig. S6.

[§]Potential residue-residue interactions (i.e. intermolecular contacts provided by Prodigy or polar contacts computed by PyMOL) at the SUMO1-S100A1 dimer interface within 5.5 Å of the 'WT, holo-NMR cluster 2' model are the following: apolar contacts (left: S100A1 dimer, right: SUMO1): B:L78/I34, A:V77/F36, B:L78/F36, A:V77/L47, B:A81/F36, B:V77/F36, B:L62/I34; salt-bridges: B:K60/E33, A:K60/E49, B:D63/R54, B:E61/K23, A:D71/K39, A:E74/K46, B:E64/R54, A:E64/K46, B:E61/K37, B:E74/R54, B:K57/E33, A:E61/K46, B:K60/D30, A:D63/K46, A:E64/K45, B:E61/R54, A:D67/H43, A:E74/H43, A:E64/H43; hydrogen bonds (bb: backbone, ss: side chain): bb A:E61/ss K46, ss A:K60/ss E49, ss A:E64/ss H43, ss A:Q73/bb K39, ss A:Q73/ss T42, bb A:V77/ss S50, ss B:K60/bb S31, bb B:L62/ss R54, bb B:E61/ss R54. The bb A:V77/ss S50 hydrogen bond was observed only for the top pose of the cluster, but not, e.g., in the top second to fourth-ranked poses (data not shown).

Supplementary Table S5: Summary of HADDOCK docking performance parameters obtained by docking SUMO1 to five cluster representatives of the holo-S100A1 dimer structure with the set II active residues.

| **S100A1 structure** | **# of clustered structures from a total of 200** | **# of HADDOCK clusters** | **Size of the best HADDOCK cluster** | **HADDOCK score of the best HADDOCK cluster*** | **HADDOCK score of the best HADDOCK cluster minus restraints violation energy multiplied by 0.1 (given in parentheses)** |
|---|---|---|---|---|---|
| WT, holo-NMR cluster 0 (representing 15 models) | 135 | 11 | 35 | -98.8 ± 6.3 | -108.2 (9.4) |
| **WT, holo-NMR cluster 1** (representing two models) | **137** | **6** | **8** | **-104.0 ± 1.3** | **-111.5 (7.5)** |
| WT, holo-NMR cluster 2 (representing one model) | 151 | 8 | 9 | -80.5 ± 12.4 | -86.3 (5.8) |
| WT, holo-NMR cluster 3 (representing one model) | 136 | 14 | 4 | -102.5 ± 18.9 | -111.3 (8.8) |
| WT, holo-NMR cluster 4 (representing one model) | 150 | 12 | 49 | -101.9 ± 4.0 | -106.3 (4.4) |

*The more negative the HADDOCK score is, the better the score is. The best scoring HADDOCK cluster is indicated by bold face. HADDOCK cluster scores are given as averages over the top-scoring four poses in the cluster, which is the default scoring approach.

Supplementary Table S6: Summary of HADDOCK docking performance parameters obtained by docking SUMO1 to five cluster representatives of the apo-S100A1 dimer structure with the set I active residues.

| **Apo-S100A1 structure** | **# of clustered structures from a total of 200** | **# of HADDOCK clusters** | **Size of the best HADDOCK cluster** | **HADDOCK score of the best HADDOCK cluster*** | **HADDOCK score of the best HADDDOCK cluster minus restraints violation energy multiplied by 0.1 (given in parentheses)** |
|---|---|---|---|---|---|
| WT, apo-NMR cluster 0 (representing 11 models) | 151 | 14 | 8 | -135.3 ± 9.6 | -146.2 (10.9) |
| WT, apo-NMR cluster 1 (representing 4 models) | 100 | 13 | 6 | -127.8 ± 9.4 | -136.7 (8.9) |
| WT, apo-NMR cluster 2 (representing 3 models) | 151 | 14 | 13 | -143.6 ± 7.3 | -153.6 (10.0) |
| WT, apo-NMR cluster 3 (representing. one model) | 148 | 14 | 6 | -142.3 ± 8.5 | -148.0 (5.7) |
| **WT, apo-NMR cluster 4** (representing one model) | **113** | **13** | **20** | **-158.2 ± 2.0** | **-171.2 (13.0)** |

*The more negative the HADDOCK score is, the better the score is. The best scoring HADDOCK cluster is indicated by bold face. HADDOCK cluster scores are given as averages over the top-scoring four poses in the cluster, which is the default scoring approach.

Supplementary Table S7: Overview of the binding affinity estimates obtained for the different SUMO1 dockings to the apo- and holo-S100A1 dimer structure cluster representatives (applying the set I active residues and always averaging over the top-scoring four poses from the best HADDOCK cluster to compute the binding affinity estimates). All values are given in kcal/mol. The structures with the best HADDOCK scores are shown in boldface. While the estimates with FoldX5.0 show that the most favorable complexes have similar affinity for apo-S100A1 and holo-S100A1, the estimates with Prodigy indicate that the complex with holo-S100A1 has a much better binding affinity than that with apo-S100A1.

| Affinity computation tool | WT, apo-NMR cluster 0 | WT, apo-NMR cluster 1 | WT, apo-NMR cluster 2 | WT, apo-NMR cluster 3 | **WT, apo-NMR cluster 4*** |
|---|---|---|---|---|---|
| Prodigy | -8.5 ± 0.7 | -7.7 ± 0.1 | -8.8 ± 0.8 | -7.9 ± 1.0 | **-7.6 ± 0.3** |
| FoldX5.0 | -4.8 ± 3.6 | -8.4 ± 2.9 | -10.6 ± 2.0 | -7.0 ± 2.4 | **-14.1 ± 1.1** |
| | WT, holo-NMR cluster 0 | WT, holo-NMR cluster 1 | **WT, holo-NMR cluster 2*** | WT, holo-NMR cluster 3 | WT, holo-NMR cluster 4 |
| Prodigy | -8.2 ± 0.5 | -7.4 ± 0.5 | **-11.2 ± 0.5** | -7.4 ± 0.8 | -8.3 ± 0.5 |
| FoldX5.0 | -8.2 ± 2.8 | -12.1 ± 2.5 | **-13.9 ± 4.3** | -6.8 ± 1.4 | -8.9 ± 3.4 |

*The more favorable binding to the holo-S100A1 is also supported by the much lower magnitude of the HADDOCK restraints violation energy for the best scored complexes for the holo-S100A1 compared to the apo-S100A1. (Table S4 vs Table S7, rows highlighted in bold). Furthermore, the docking for the SUMO1-S100A1 apo-NMR cluster 4 shows the highest value for the restraints violation energy among the dockings with apo-S100A1 (Table S7), while the docking for the SUMO1-S100A1 holo-NMR cluster 2 has a restraints violation energy among the lowest values within the set of dockings to holo-S100A1 (Table S4). This indicates lower consistency between the experimental restraints and a good docking score for the apo-S100A1 than for the holo-S100A1.

Supplementary Table S 8: Results of SDA docking. Clustered encounter complexes are ordered by SDA-cluster size. According to the SDA-cluster sizes, SUMO1 approaches the S100A1 dimer preferentially by facing the “arched cleft” (defined here as from “above”) (“above”, red rows: 17406384 recorded encounter complexes; “below”, blue rows: 55362 recorded encounter complexes) (see Fig S10-C for an overview of the relative positions of all SDA-clusters). The representatives of SDA-clusters 5 and 6 (rows in bold face), which correspond to the symmetric HADDOCK binding modes, have the lowest total interaction energies. SDA-cluster 5 corresponds to the second-best and SDA-cluster 6 to the best HADDOCK cluster of the SUMO1 docking to ‘WT, holo-NMR cluster 2’ applying the set I active residues. The SDA-clusters 1 and 2 are more populated and have similar average interaction energies to SDA-clusters 5 and 6, but they have a much higher structural variability (i.e., spread and standard deviation). Energies are given in kcal/mol.

| SDA-cluster id | SDA-cluster size | Interaction energy of SDA-cluster representative | Electrostatic interaction | Electrostatic desolvation | Non-polar desolvation | Average interaction energy of SDA-cluster | Spread | Standard deviation |
|---|---|---|---|---|---|---|---|---|
| 1 | 8637146 | -26.63 | -13.92 | 6.67 | -19.39 | -31.50 | 12.35 | 4.07 |
| 2 | 5628399 | -28.15 | -13.89 | 6.92 | -21.18 | -30.50 | 12.23 | 4.31 |
| **5** | **2274838** | **-29.73** | **-15.54** | **6.36** | **-20.55** | **-30.93** | **0.81** | **0.32** |
| **6** | **825849** | **-29.99** | **-15.62** | **6.46** | **-20.84** | **-30.49** | **0.76** | **0.46** |
| 4 | 43541 | -27.17 | -12.01 | 4.93 | -20.09 | -26.98 | 1.24 | 0.73 |
| 3 | 24830 | -25.63 | -15.10 | 5.92 | -16.45 | -26.96 | 8.02 | 3.32 |
| 7 | 15322 | -26.76 | -15.29 | 6.06 | -17.54 | -28.81 | 5.16 | 2.41 |
| 8 | 10939 | -26.87 | -11.55 | 4.87 | -20.19 | -27.48 | 1.73 | 2.22 |
| 9 | 625 | -25.08 | -10.54 | 4.49 | -19.02 | -24.85 | 0.82 | 0.28 |
| 10 | 257 | -26.19 | -11.84 | 5.48 | -19.82 | -26.12 | 0.53 | 0.45 |

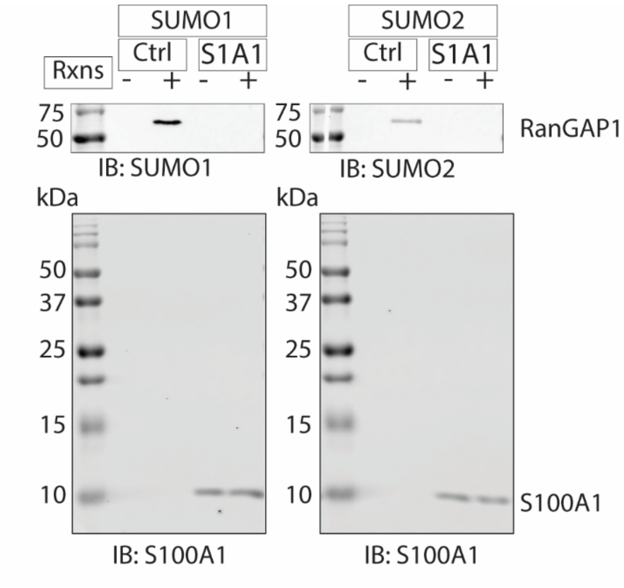


Supporting Figure-1: In vitro S100A1 SUMOylation assay. The assay was performed under original/ kit assay conditions. Western blotting analysis of *in vitro* SUMOylation assay (left: SUMO1; right: SUMO2) conducted at 37°C for 1 hr in the indicated conditions using antibodies directed against SUMO1, SUMO2 and S100A1. The assay control RanGAP1 is SUMOylated, while S100A1 is not SUMOylated.

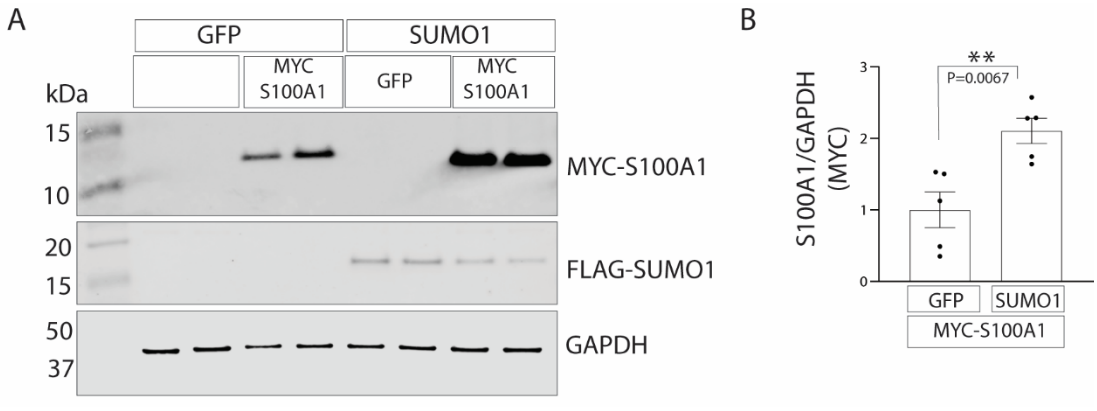


Supporting Figure-2: S100A1 and SUMO1 co-(over) expression increases S100A1 protein abundance in COS1.

A) MYC-tagged S100A1 was co-overexpressed together with either GFP or SUMO1 in COS1 using adenoviral gene delivery. 24 hrs post COS1 adenoviral transduction, cells were processed for western blotting analysis and probed with antibodies directed against MYC, FLAG, and GAPDH.

B) Quantification of S100A1 protein fold change normalized to GAPDH co-(over) expressed together with either GFP or SUMO1. Values are presented as mean ± s.e.m., n=5 (independent samples).

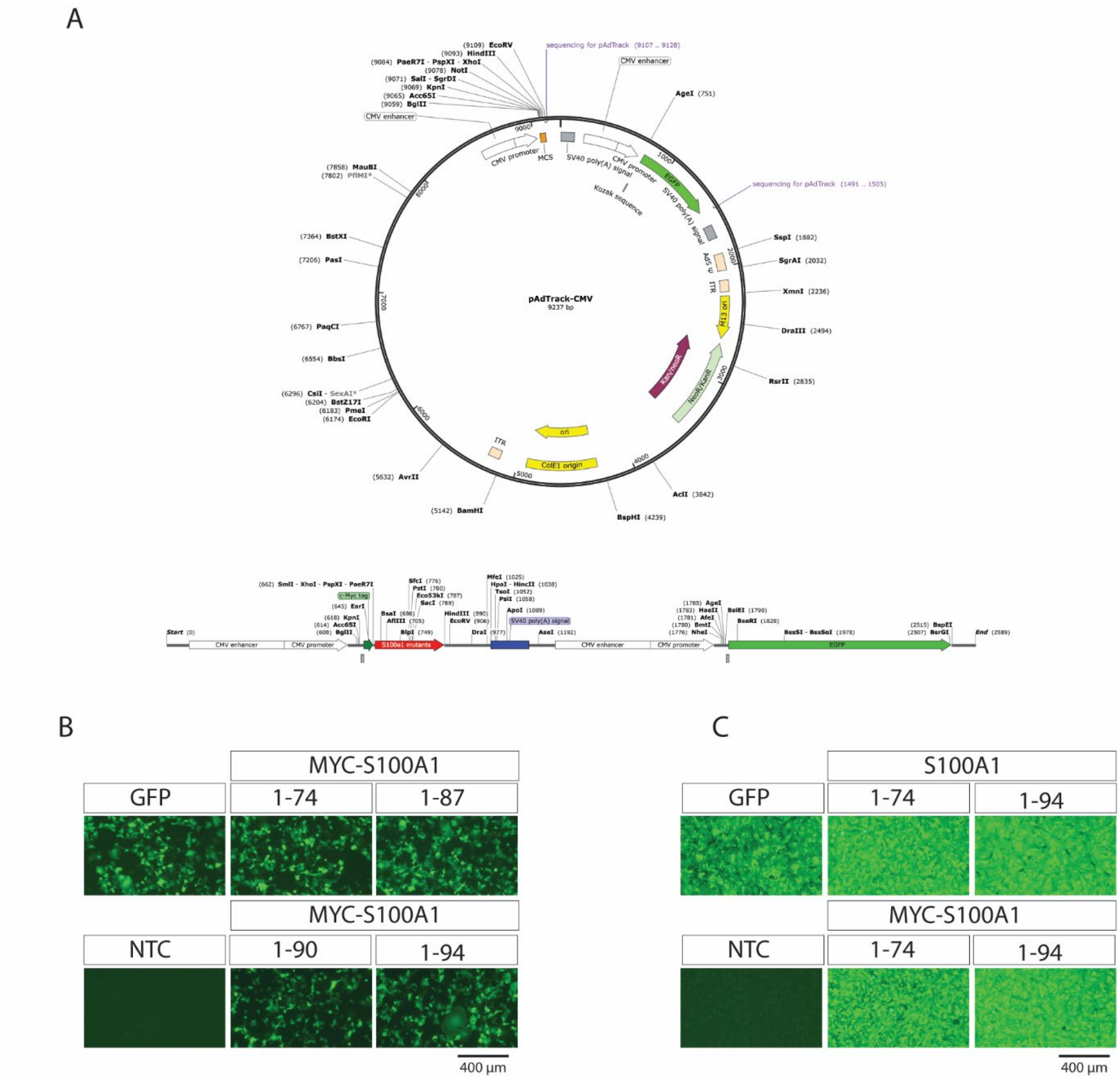


Supporting Figure-3: Expression vector map and transfection and transduction control EGFP signal.

A) pAdTrack-CMV vector map (top) and S100A1 and S100A1 mutants cloning sites on pAdTrack-CMV vector (bottom).

B) EGFP fluorescence signal of COS1 cells transfected with indicated plasmids.

C) EGFP fluorescence signal of NRVM transduced with indicated adenovirus constructs.

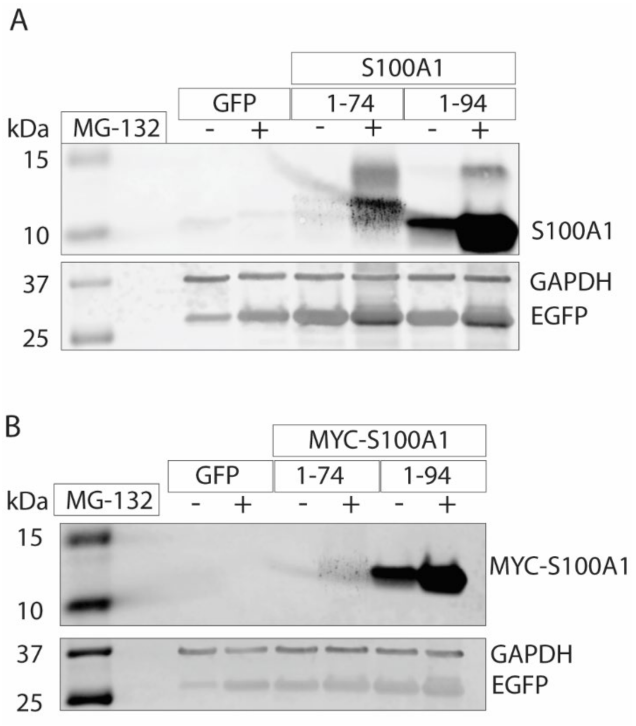


Supporting Figure-4: Proteasome inhibition blocked degradation of the S100A1 truncation mutant S100A1-1-74.

A) Western blotting analysis of COS1 cells transduced with untagged S100A1-1-74 and S100A1-1-94 and the control EGFP in the absence or presence of 10 µM MG-132 (a proteasome inhibitor). 24 hrs post transduction, samples were processed for western blotting and probed with antibodies directed against S100A1, GAPDH and GFP. Western blotting analysis revealed that S100A1-1-74 could be detected at the protein level in the presence of 10 µM MG-132

B) Western blotting analysis of COS1 cells transduced with MYC-tagged S100A1 (1-74), MYC-tagged S100A1 (1-94); and the control EGFP in the absence or presence of 10 µM MG-132 (proteasome inhibitor). 24 hrs post transduction, samples were processed for western blotting and probed with antibodies directed against S100A1, GAPDH and GFP. COS1 cells were treated with either DMSO (-) or 10 µM MG-132 (+) for 10 hrs prior to harvest. Western blotting analysis revealed that MYC-tagged S100A1 mutants; 1-74 (devoid of SIM) could be detected at the protein level in the presence of 10 µM MG-132.

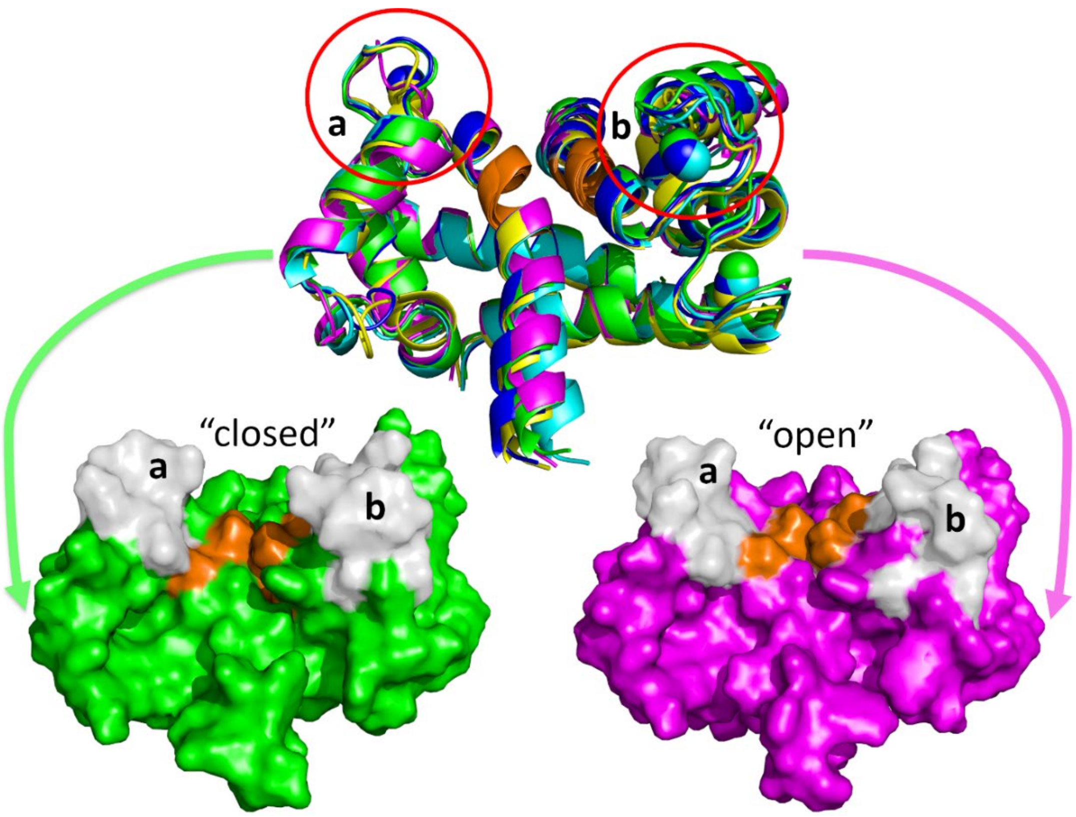


Supporting Figure-5: Flexibility within the loops of the C-terminal EF-hand motifs (highlighted via circles and labeled a and b) of S100A1 affects the accessibility of the SIMs. The core residues (i.e., $_{76}$VVLVA$_{80}$) of the two SIMs in the S100A1 homodimer are highlighted in orange in structures from the NMR structure of the holo-S100A1 homodimer (PDB id 2LP3). Top: cartoon representation of the structures representing conformational clusters (cluster 0: green, cluster 1: cyan, cluster 2: magenta, cluster 3: yellow, cluster 4: blue). Calcium ions are shown as spheres. Bottom: surface representations of the S100A1 dimer structure in cluster 0 (green, NMR model 1, closed) and cluster 2 (pink, NMR model 9, open). The respective loops a and b are highlighted in grey.

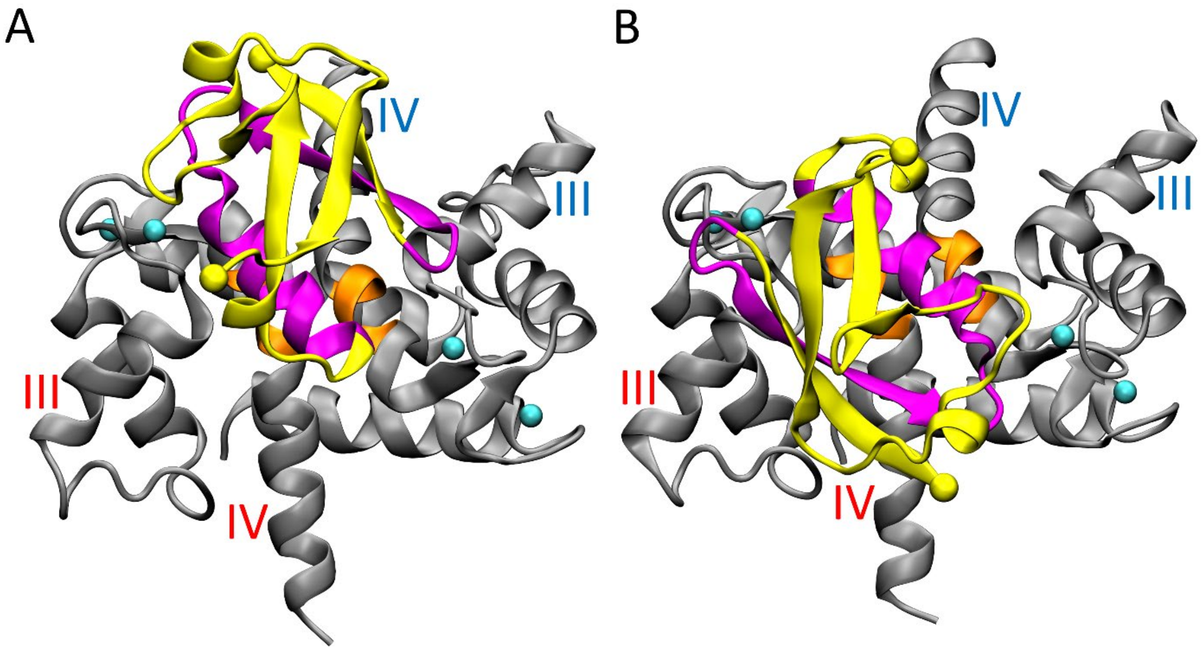


Supporting Figure-6: The two symmetric SUMO1-S100A1 homodimer (i.e., 'WT, holo-NMR cluster 2') binding modes predicted by docking with HADDOCK. SUMO1-S100A1 dimer complexes are color coded according to Figure 6. The S100A1 helices III and IV are highlighted in red (chain A) or blue (chain B).

A) Top view of the SUMO1-S100A1 dimer binding mode (best-scoring pose of the best cluster).

B) Top view of the symmetric binding mode obtained for the best-scoring pose of the second cluster obtained from docking SUMO1 to the S100A1 dimer.

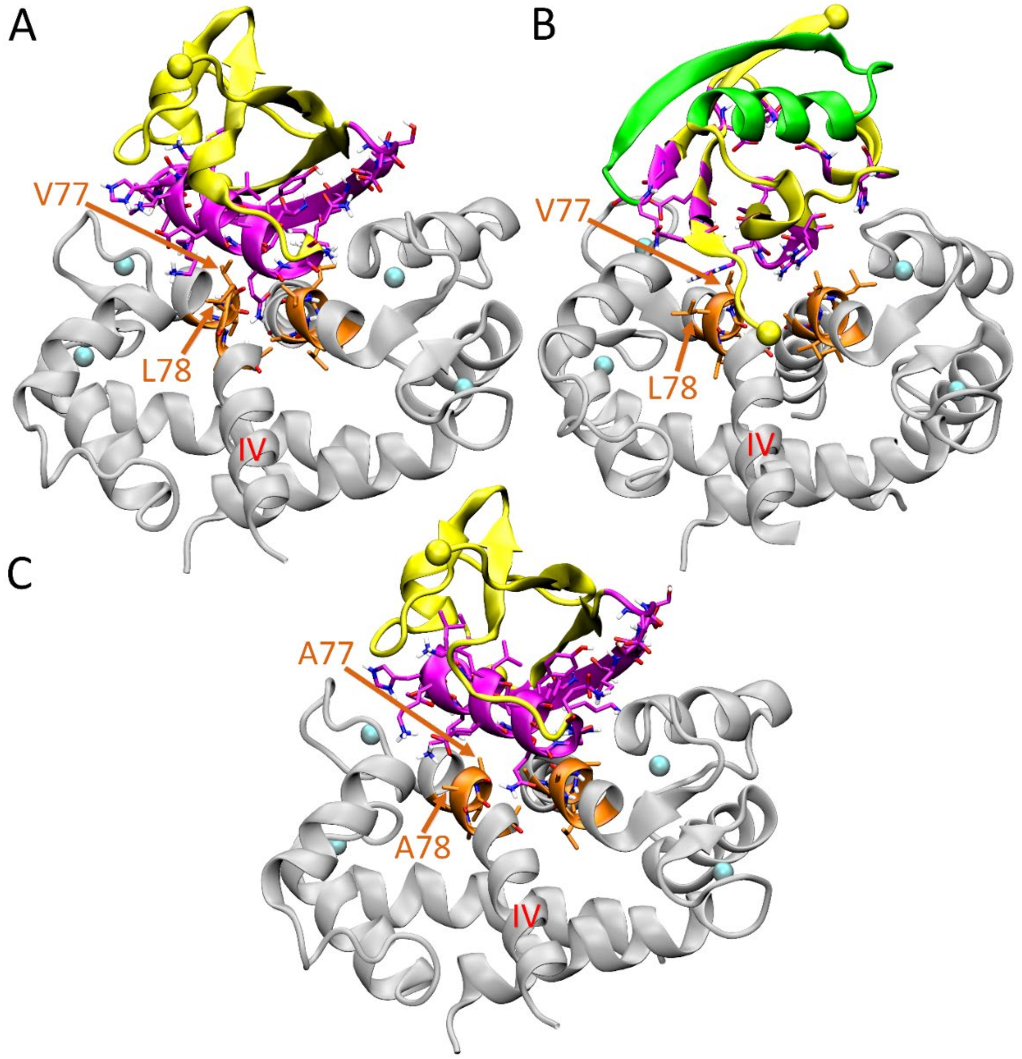


Supporting Figure-7: HADDOCK-predicted top ranked binding modes of SUMO1 in complex with the S100A1 dimer.

A) SUMO1 S100A1-WT dimer complex, applying the set I active residues.

B) SUMO1 S100A1-WT dimer complex, applying the alternative set II active residues.

C) SUMO1 S100A1-AAA dimer complex, applying the set I active residues. The SUMO1-S100A1 dimer complex is color coded according to Figure 6, except that the area corresponding to the set II active residues is highlighted in magenta (SUMO1 residues 28, 29, 63, 66-70, 75, 81, 83, 86, 89, 91) in subpanel B, whereas the area corresponding to the set I active residues is shown in green in subpanel B. All residues that were defined as active residues are highlighted in stick representation. For orientation, one of the helices IV is labelled in red, while the residues 77 and 78 in the corresponding SIM are labelled in orange.

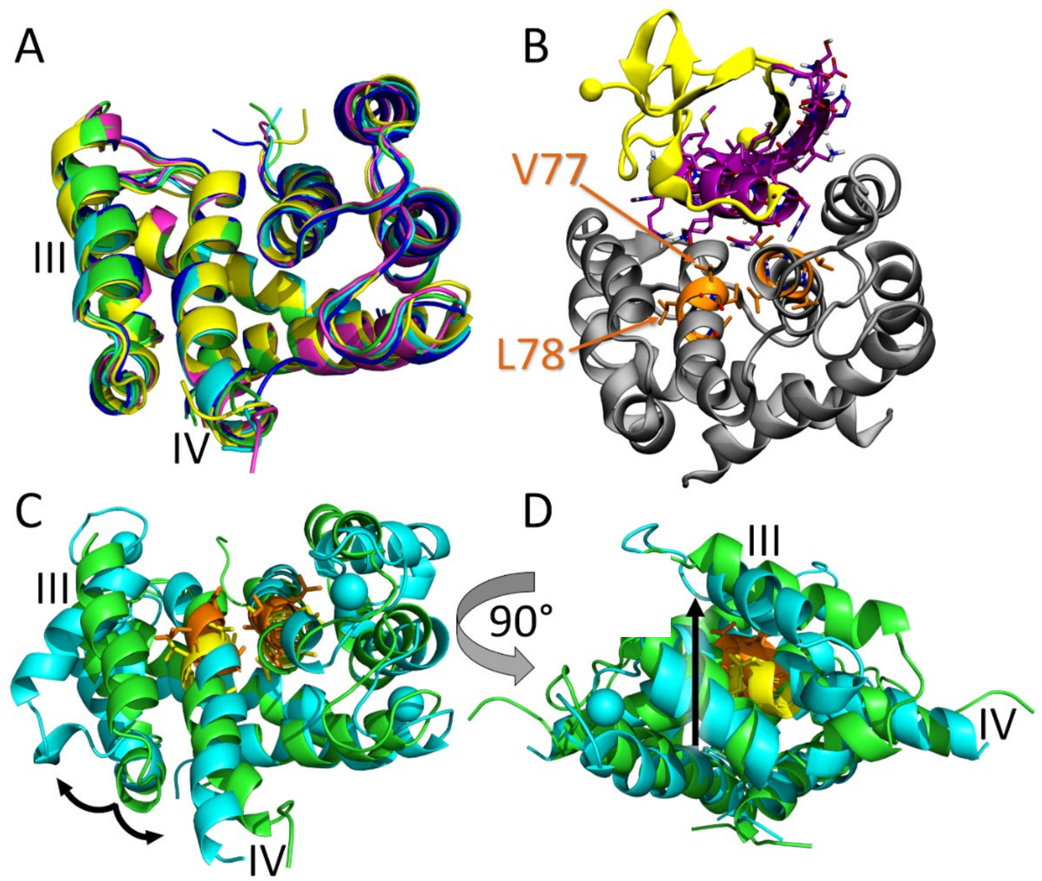


Supporting Figure-8: Docking of SUMO1 to the apo-S100A1 dimer structure indicates that SUMO1-SIM interaction might be constricted by the apo-S100A1 dimer structure. Helices III and IV are indicated for one of the S100A1 subunits.

A) Cartoon representation of the apo-S100A1 NMR structure (PDB id 2L0P) conformational clusters (cluster 0: green, cluster 1: cyan, cluster 2: magenta, cluster 3: yellow, cluster 4: blue). Clustering was performed analogously to the holo-S100A1 dimer structure. Contrary to the holo-S100A1, there was no striking conformational variability visible between the different cluster representatives.

B) Best-scoring pose between SUMO1 and the apo-S100A1 homodimer (apo-NMR cluster 4). Color coding is according to Figure 6. All residues that were included as active residues are highlighted in stick representation. The residues 77 and 78 in the corresponding SIM are labelled in orange. The overall pose is similar to those obtained for holo-S100A1, which is not surprising given the restraints for the interaction with the SIM.

C+D) Comparison between the holo- and apo-structures of the S100A1 homodimer. The holo-structure is shown in cyan (2LP3, model 1), the apo-structure (2L0P, model 1) is depicted in green. A front view (C) and a side view (D) of the superimposed holo- and apo-S100A1 homodimer structures is depicted. The SIM is highlighted in orange or yellow, respectively, with SIM residues shown as sticks and an arrow (right) highlighting the different positioning of the SIM between apo- and holo-S100A1 structures. The calcium ions in the holo-structure are shown as spheres. Helices III and IV (and their repositioning between the holo and apo states) are indicated for one of the S100A1 subunits.

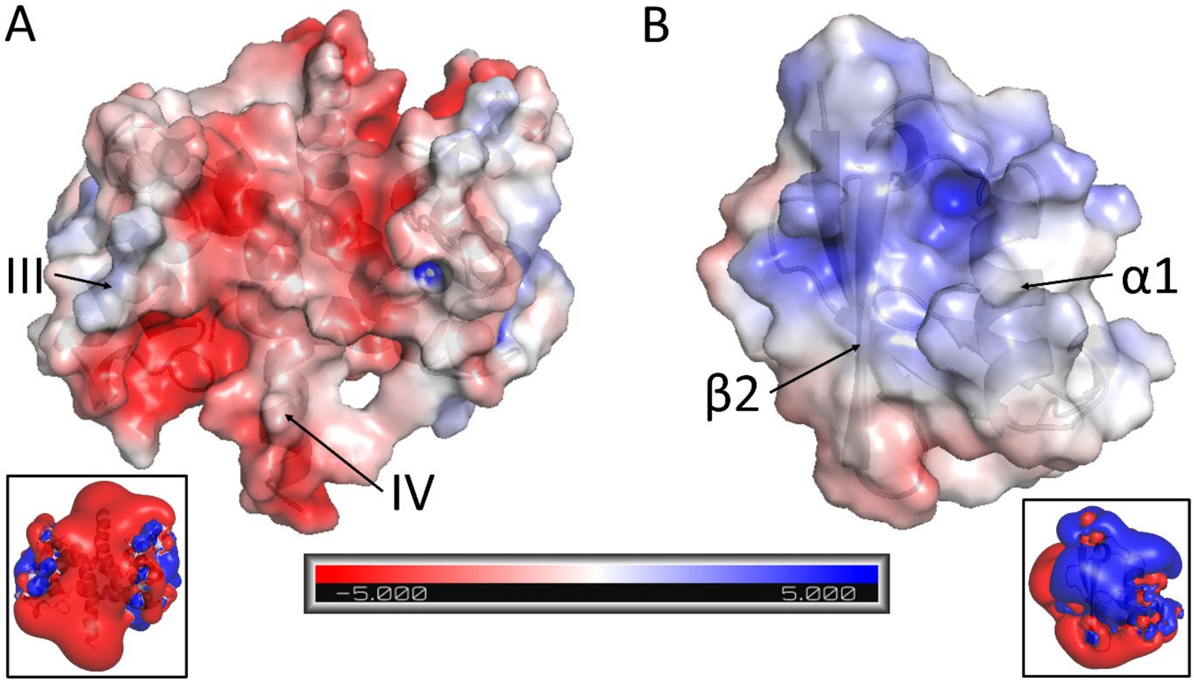


Supporting Figure-9: The electrostatic potentials of the predicted (set I) holo-S100A1 dimer ('holo-NMR cluster 2') (A) and SUMO1 (B) docking interfaces are complementary. The proteins are displayed as transparent molecular surfaces with electrostatic potentials projected onto them and grey cartoon representations (including grey spheres for the $Ca^{2+}$ ions of holo-S100A1). H-III and H-IV of S100A1 as well as the α1-helix and the β2-strand of SUMO1 are labeled. Negatively charged regions are shown in red, positively charged regions in blue, and neutral regions in white. The color bar shows the respective range from -5.0 to +5.0 kT/e. The insets show the respective ± 1.0 kT/e electrostatic potential isocontours of the proteins in the same orientation.

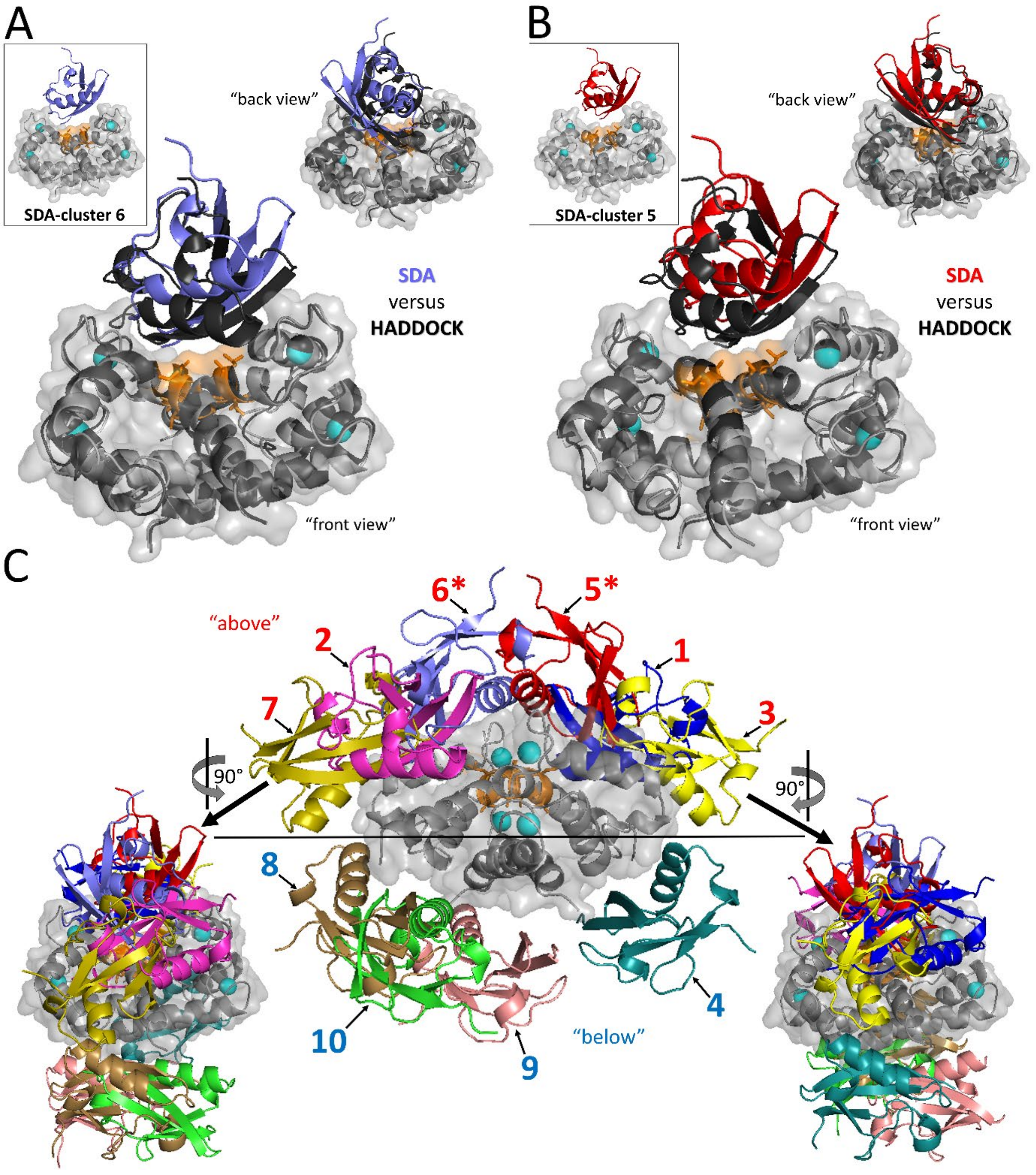


Supporting Figure-10: Docking of SUMO1 to the holo-S100A1 dimer ('holo-NMR cluster 2') with SDA predicts diffusional encounter complexes that are overall similar to the respective bound complexes predicted by HADDOCK. The holo-S100A1 dimer structure 'holo-NMR cluster 2' (whose conformation is treated as rigid by SDA) is depicted in grey cartoon representation with a transparent molecular surface and with $Ca^{2+}$ ions shown as cyan spheres and the SIM residues 'VVLVA' highlighted in orange stick representation. SUMO1 is shown in cartoon representations of different colors in the 10 encounter complexes obtained with SDA. (A,B) Encounter complex representatives for SDA-cluster 6 (A, violet) and 5 (B, red). The most similar binding modes predicted by HADDOCK (i.e., best-ranking poses of the best-scoring and second best-scoring HADDOCK cluster of SUMO1 docking to 'WT, holo-NMR cluster 2' applying the set I active residues) are shown in black cartoon/spheres representation for comparison. The insets show views from the "back". C) Overview of all ten SDA-clusters obtained with the holo-S100A1 dimer structure depicted from the side (i.e., a perpendicular view towards the 'arched cleft'). The SDA-clusters similar to HADDOCK-predicted binding modes (i.e., SDA-cluster 5 and 6) are additionally labeled with an asterisk. SUMO1 poses that approach the holo-S100A1 dimer from “above” (i.e., facing the ‘arched cleft’) have red labels, while those approaching from “below” have blue labels. SUMO1 docking encounters approaching the holo-S100A1 dimer from “above” significantly outweigh those approaching from “below” (see Table S8). The left and right insets show “front” and “back” views of the SDA-clusters docking to the holo-S100A1 dimer. Note that no SDA-clusters corresponding to SUMO1 encountering the holo-S100A1 dimer from the side were identified.

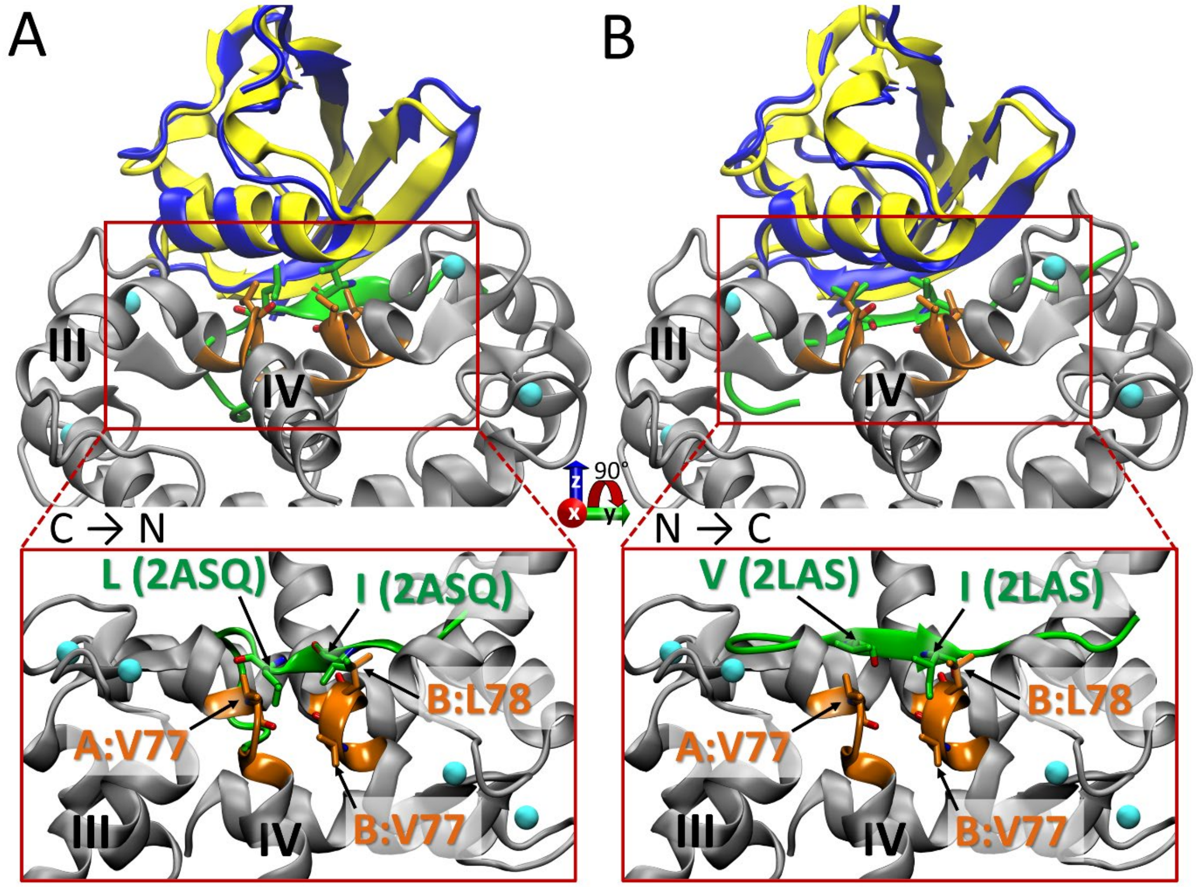


Supporting Figure-11: Comparison between the docked SUMO1 S100A1-WT homodimer binding mode (top ranked pose) and "classical" interactions of SIM-containing peptides with SUMO1. The coloring and representation of the SUMO1-wild-type S100A1 homodimer complex is the same as in Figure 6. Helices III and IV are indicated for one of the S100A1 subunits.

Comparison to the NMR structures 2ASQ (A) and 2LAS (B). SUMO1 from 2ASQ or 2LAS, respectively, is shown in blue cartoon representation, superimposed on SUMO1 (all yellow) in the docked SUMO1-S100A1 complex. The bound peptide (KVDV**I**D**L**TIESSSD or DNEIE**V**I**I**VWEKK, respectively) is shown as green cartoon with the apolar peptide residues occupying the hydrophobic cleft of SUMO1 (highlighted in bold type in the sequences depicted above) shown as sticks (the two peptides bind SUMO1 with oppositely aligned termini, indicated via C→N or N→C, respectively). The S100A1 dimer residues A:V77, B:V77, and B:L78 are also shown as sticks. Besides the side view, an approximately perpendicular top view without SUMO1 structures is shown in the inlays below. S100A1 and peptide residues highlighted in stick representation are additionally labelled in orange or green, respectively, in the inlays below.
